# Neural spiketrains and population vectors entangle neural representations

**DOI:** 10.64898/2026.08.27.744867

**Authors:** Melvin Vaupel, Valdemar Kargård Olsen, Sigurd Gaukstad, Erik Hermansen, Benjamin A. Dunn

**Affiliations:** Department of Mathematical Sciences, NTNU, Trondheim, Norway; Kavli Institute for Systems Neuroscience, NTNU, Trondheim, Norway

## Abstract

Neural recordings are usually analyzed by comparing neural spiketrains or comparing time bins (population vectors). If multiple variables drive the neural activity these comparisons will be affected by all of them. Our aim is to disentangle the different latent variables or covariates that drive neural activity and reveal their structure and geometry. The central idea of the paper is that a matrix is disentangled when its rows and columns are local on each other, a condition we call bidirectional locality. In such a matrix, rows and columns encode the same geometry and they respond to only one localized part of it. This suggests finding bidirectional local matrices in a given data matrix, from which we can recover the geometry of the covariates driving it in a straightforward way. We present two ways of doing just this. The first method, coherent projections, works by finding non-negative projections of the neural data matrix (neurons by time bins) that are bidirectionally local. The second method, clumps, works by finding dense submatrices of the neural data matrix, that each identify a local region of one covariate. We apply these methods to two neural datasets, showing that they can separate grid cell modules and reveal a movement-driven low-dimensional structure in the motor cortex.

## 1 Introduction

During most of our waking hours, we perceive compositions of many unrelated things. The experience of reading this paper is composed of the font size of the text, the brightness in the environment, your posture, your mood, your previous knowledge of the topic, and so on. All these variables can be combined in many ways and it is ultimately these combinations that make up your experience. Moreover, the number of possible such combinations increases exponentially as we add more variables. For animals and machines to be able to learn about and represent this complex world in an efficient way, it is necessary to break it down into smaller, low-dimensional parts^1–10^. We can do this because each low-dimensional part functions as an independent source of variation, which we refer to as a *covariate*, regardless of whether or not it is directly observable.

In a similar way, neural data that contains hundreds to thousands of neurons simultaneously recorded, can include neurons from multiple functional circuits tuned to multiple covariates^11–19^. It is often difficult to make sense of such complex and apparently high-dimensional datasets. Like above, breaking these compositions up into several low-dimensional parts makes it easier for us to understand them. Another way of saying this is that we want to disentangle the representations^20^ of several covariates entangled in the data. This is a challenging problem in general^17;21;22^, but even more so in an unsupervised setting^2;23–25^, where we do not have access to the covariates themselves.

Traditional ways of analyzing large neural datasets, such as dimensionality reduction and clustering, rely on comparing either neurons (the co-activity perspective) or time bins (the population vector perspective). This implicitly assumes that neurons or time bins are point-like on the covariate we want to find. In the population vector view, time bins are collapsed into points, entangling covariates that are present simultaneously. Similarly, methods based on neural co-activations entangle neurons that respond to several covariates. We show a schematic version of this argument in Figure 1, where the data contains a population of neurons tuned to a ring and another tuned to three states. We see that treating time bins as points entangles these two populations, giving us the product of a circle and three states (i.e., three circles) in the column space (1c). Moreover, the number of unique population vectors grows exponentially as we record data tuned to more covariates (1d). We suspect that this exponential growth in the number of possible population vectors is part of the reason why large recordings look complex and high-dimensional.

**Figure 1:**
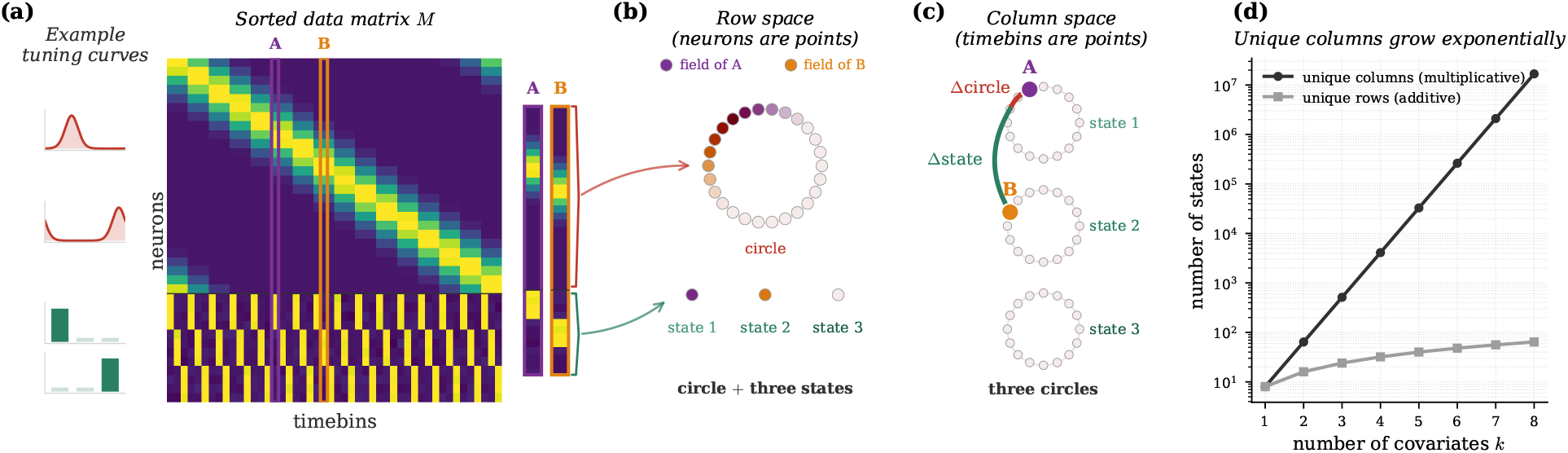
Neurons and time bins are often not point-like on a single covariate. **(a)** In this example, we use two independent covariates; one that is circular and the other describing three discrete states. This choice of covariates show that the principles we discuss here hold for both discrete and continuous covariates. We have one population of neurons tuned (von Mises) to the circle and another tuned to one of the three states, as shown in the example tuning curves to the left. Both the neurons and time bins of the data matrix are sorted according to the covariates, making its structure clear. We pick two probe columns (or time bins or population vectors) that we follow through panel B and C. **(b)** Making every neuron a point, we see that they form one continuous circle and three discrete states, mirroring their local tuning to one covariate. The neurons in the row space are colored by the activity of the two probe columns; their intrinsic field. We see that the two columns are each active at one local part of the circle and one of the three states, mirroring the covariate states at that column or time bin. **(c)** Making every time bin a point, we see that they form three disjoint circles. This is the Cartesian product of a circle and three discrete points. The two probe columns are marked in the column space. We can see that the distance between them depends on both their distance on the circle (Δ**circle**) and their distance on the three states (Δ**state**). This reflects the fact that any scalar comparison of the two probe columns, will be affected by the entire column — both the upper and lower part of the vectors displayed to the right in panel (a). **(c)** This shows how the number of rows and columns grows with the number of covariates (in this case, with 8 states each). The number of unique columns grows multiplicatively, giving exponential growth, while the number of unique rows grows only additively. There is of course nothing special about rows vs. columns here. When the covariates are entangled in the data, in rows or columns, it quickly looks complicated.

### More concisely

*In typical large population recordings of neurons tuned to multiple covariates, both the neural co-activities and the population vectors entangle the geometric structure of those covariates*.

In this paper, we propose a novel perspective on the neural disentanglement problem that revolves around the concept of *locality* and in particular the idea that the activity matrix of a disentangled representation carries a signature that we call *bidirectional locality*. To formalize this signature we introduce the notion of *intrinsic fields*. These allow us to study what individual neurons look like on the space of all time bins (population vectors) and dually, what individual time bins look like on the space of all neurons (co-activation vectors).

We will argue that bidirectional locality provides us with a tool to test whether the point-like assumption holds for rows and columns simultaneously. Importantly, bidirectional locality should be interpreted as one joint condition. The locality of rows on the column space is only meaningful when the column space is itself a faithful representation of the covariates, and likewise in the other direction — so the two requirements constrain each other and cannot be evaluated separately. Fortunately, this condition can be checked on the data matrix directly, without reference to the (observed or unobserved) covariates. This makes it useful both as a diagnostic and as an optimization target.

We introduce two methods that use bidirectional locality to find representations of different covariates. The first, *coherent projections*, learns projections of the data matrix that are explicitly optimized for bidirectional locality. By learning multiple such projections and encouraging them to be decorrelated, an entangled recording is separated into one matrix per covariate, each disentangled enough for standard analyses to recover the underlying covariate. The second method scans the data matrix for dense submatrices, defined by ensembles of neurons collectively active on a subset of time bins. Our intuition is that such coactivity is driven by one small region of a single covariate, making both the rows and columns selected by the submatrix local on that covariate. These dense submatrices, which in reference to Good^26^ we call *clumps*, are good candidates for points in a space that represents the underlying covariates. We validate both methods on synthetic data and several neural datasets. In particular, we separate grid cell modules (Section 2.6) and show how running and turning speed are associated with a ring-like structure in the primary motor cortex (Section 2.7). This illustrates that bidirectional locality can be an important tool for revealing how the brain breaks up the world into low-dimensional parts.

## 2 Results

### 2.1 Intrinsic fields and bidirectional locality

Consider the activity of *N* neurons recorded over *T* time bins. We begin with the simplest case, every neuron tuned to a single covariate *X*, and build up from there. Neuron *i* responds to the state *x*_*j*_ present at time bin *j* through a tuning function *f*_*i*_, giving activity *f*_*i*_(*x*_*j*_). We collect these entries into an *N* × *T* data matrix

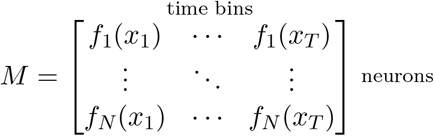

Defined in this way, a covariate is not any arbitrary recorded variable — it is what the neurons (and time bins) are ultimately tuned to. Most standard analyses summarize a data matrix using similarities or distances between either time bins^27–29^ or neurons^30–34^, and use the resulting space to infer or decode the geometry of the underlying covariate. This is the population vector and co-activation perspective on *M*, respectively. Each column of *M* is the activity of all neurons at a single time bin. In the population vector view each time bin gets compressed into a single point and compared through their similarities and dissimilarities across all neurons. Dually, each row of *M* is one neuron’s activity across the whole recording. In the co-activation view we compress these into points, and distances relate entire neural spike trains over all time. Almost every population analysis adopts one view or the other, and with it an implicit, rarely-questioned assumption: that the entities treated as points (time bins in one view, neurons in the other) are *point-like* on the underlying covariate, each occupying a single small and local region. We argue that this assumption can and should be tested, not silently assumed. The synthetic examples that follow show the consequences of ignoring it.

#### Intrinsic fields

Given a row *i* in data matrix *M*, we can identify which points it activates in the space of population vectors (columns). We call this the *intrinsic field* of the row. Formally, it is just *j* 1→ *M*_*ij*_, the row itself viewed as a function on the columns. Alternatively, we could also normalize it to a probability distribution (Methods, § 4.1.2). Dually, the intrinsic field of column *j* on the row space is *i* 1→ *M*_*ij*_. Both fields are defined from *M* alone, with no reference to any external covariate, and they are exactly what we visualize when we color an embedding of columns by a probe row *i*_0_, or an embedding of rows by a probe column *j*_0_ (see Figure 2).

**Figure 2:**
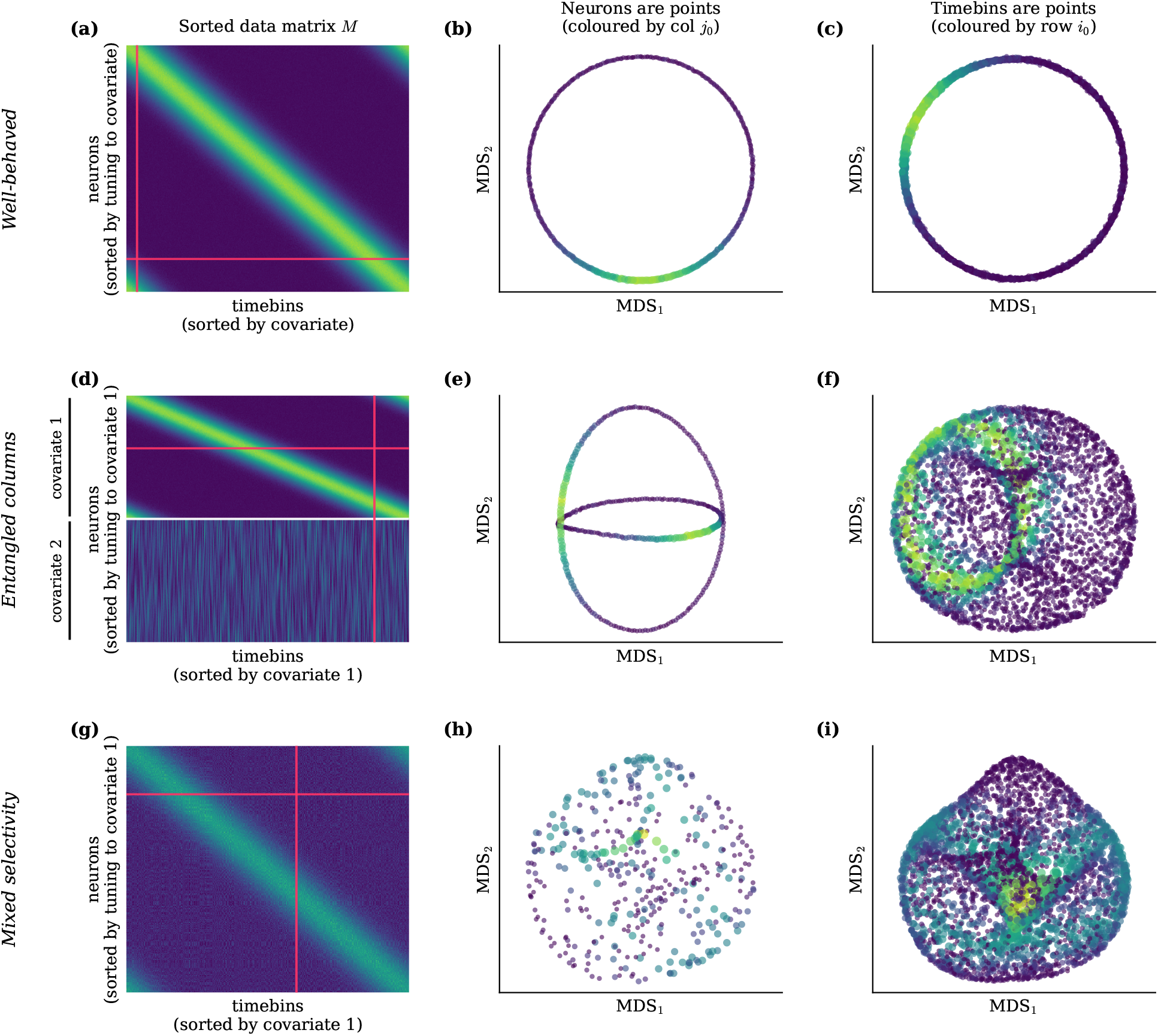
Bidirectional locality is the signature of a disentangled representation. Three synthetic cases on continuous data (*X* = *S*^1^ circuits, von Mises tuning *κ* = 4). Each row is one case; columns are the data matrix *M*, the row-MDS (neurons, each colored and scaled by a probe column *j*_0_) and the col-MDS (time bins, each colored and scaled by a probe row *i*_0_). Red lines on *M* mark the two probes. **(a–c)** Well-behaved: *N* = 400 neurons tuned to a single circle over *T* = 4000 time bins. Both embeddings recover the circle and each probe lights a single localized region. Bidirectional locality holds. **(d–f)** Entangled columns: *N* = 400 neurons tuned to one of two circles over *T* = 4000 time bins. The row-MDS recovers the disjoint union *X*_1_ ⊔ *X*_2_ (two circles), while the col-MDS recovers the product (torus-like cloud). The intrinsic field of the probe neuron is local on the circle it is tuned to but wraps around the other circle. Locality fails on the column side. **(g–i)** Mixed selectivity: *N* = 400 neurons tuned to two circles over *T* = 4000 time bins. Both embeddings show the entangled product geometry and both intrinsic fields are non-local, looking like two circles connected at a point (like in Extended Data Figure 8a). Locality fails on both sides. The discrete-data version is in Extended Data Figure 7.

### 2.2 Bidirectional locality is the signature of a disentangled representation

#### A single covariate

Figure 2a–c shows the well-behaved case, where every neuron has unimodal, or local, tuning to a single covariate. We use a circle, *S*^1^, for simplicity, but the covariate can have any shape, discrete or continuous. We embed both the rows (b) and columns (c) of the data matrix *M* into two dimensions, using Multidimensional Scaling (MDS). Note that there is no reason for using MDS over any other dimensionality reduction method, other than it preserving the covariate geometry well for illustration. Both the row-MDS and the col-MDS recover the geometry of *X* (i.e., a circle). Moreover, a probe row *i*_0_ lights up a single localized region on the col-MDS, and dually a probe column *j*_0_ on the row-MDS. These localized intrinsic fields reflect the locality of neurons and time bins on the underlying covariate, which justifies representing them as points in their MDS embeddings in the first place. In the rest of this section we study several situations in which circular covariates are entangled and the bidirectional locality breaks, using the intrinsic fields on MDS embeddings as a visual diagnostic. In the Extended Data, we show analogous situations for discrete objects whose properties (e.g., shape and color) are entangled in a data matrix. This highlights that bidirectional locality is a useful diagnostic not only for continuous manifolds, but also for discrete data.

#### Multiple neural populations entangle population vectors

Modern neural recordings can contain thousands of neurons recorded at once. It is unlikely that all of these neurons happen to be tuned to the same covariate. Assume that we have recorded neurons tuned to *K* different covariate spaces *X*_1_, …, *X*_*K*_. At each time bin we sample a combination of states: one from each covariate. A time bin *j* can thus be described with a tuple (*x*_*j*1_, …, *x*_*jK*_). Formally speaking, these are samples from the product space of the covariates *X*_1_ × … × *X*_*K*_. We write the tuning of neuron *i* to covariate *k* at time bin *j* as *f*_*ik*_(*x*_*jk*_). In this example we assume that every neuron is tuned only to one of the covariates. This makes it possible to write the data matrix *M*_*ij*_ = ∑_*k*_ *f*_*ik*_(*x*_*jk*_) as a stack of smaller data matrices *M*_*k*_, containing the neural population tuned to covariate *k*:

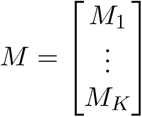

Figure 2d–f shows the population vector and co-activation view on such a matrix, for circular covariates (the discrete case is shown in the Extended Data). Each neuron is still locally tuned to a single covariate, so the row-MDS recovers the disjoint union of the two covariate spaces (two rings). A time bin, in contrast, is a point on both covariate spaces at once, carrying a state (*x*_*j*1_, *x*_*j*2_) on each circle. Its intrinsic field on the row space reflects this: rather than a single localized spot, it lights up a localized region on each ring. Accordingly, the col-MDS of the non-local time bins recovers the product *X*_1_ × *X*_2_ as a torus-like cloud, not two circles. A single neuron is now local on the covariate it is tuned to and spreads over the entire covariate it is not tuned to, resulting in a circular intrinsic field. Because the number of combinations of states from different covariates grows exponentially with the number of covariates, the product is in general only sparsely sampled, which can make the time bin space look disorganized in real data. In short, when stacking distinct populations of local neurons, the neurons remain local but the time bins do not. Because the neurons are still local on the covariates, the issues in this example could be resolved by clustering the neurons and looking at each cluster separately. As we will see next, this strategy does not work in general.

#### Mixed selectivity entangles co-activations and population vectors

A typical phenomenon in neural coding is *mixed selectivity*, where single neurons are tuned to more than one covariate^11;16–18^. Like in the previous section, we can model this with a data matrix generated as *M*_*ij*_ = ∑_*k*_ *f*_*ik*_(*x*_*jk*_). The difference is that now the neurons can be tuned to more than one covariate: for each neuron *i*, multiple of the *f*_*ik*_s can encode information. In particular, the matrix no longer admits its stacked form from above. Note that neurons would also be entangled if they responded differently to the covariates at different times, perhaps during different parts of a task^35^.

Figure 2g–i shows the continuous example with mixed tuning to two circles (again, the discrete version is found in the Extended Data). Now, every neuron is tuned to both covariates, so both the row-MDS and the col-MDS recover the same product geometry. Despite the rows and columns agreeing, neither side is local on the other: a neuron fires whenever its tuning matches either the state of circle 1 or circle 2 (additive mixed selectivity; see below). Dually, in a time bin, states from both covariates are present simultaneously. The intrinsic fields of neurons and time bins are therefore both non-local and, with sufficient sampling, should resemble two circles glued together at a point.

#### Bidirectional locality

Across these three cases, only the disentangled one (Figure 2a-c) is bidirectionally local: rows and columns are both point-like and intrinsic fields are localized in both directions. With multiple pure populations (Figure 2d-f), locality fails on the column side as time bins activate on both covariates simultaneously. Under mixed selectivity (Figure 2g-i) it fails from both perspectives. This signature is intrinsic to the data and can be read from *M* alone, with no reference to the underlying covariate.

The bidirectional nature of the test is essential. In the entangled-columns case a neuron is unimodally tuned to a single covariate, yet its intrinsic field on the column space is still spread out (a circle), because that space is itself the product *X*_1_ × *X*_2_, not either component alone. Thus, the shape of an intrinsic field can be no more informative than the space it is measured in. The locality of rows on the column space is meaningful only when the columns are themselves locally organized, and likewise in the other direction; bidirectional locality is the joint condition that closes this loop.

### 2.3 Additive versus multiplicative mixed selectivity

The mixed-selectivity model *M*_*ij*_ = ∑_*k*_ *f*_*ik*_(*x*_*jk*_) above is *additive* (OR-like): the neuron activates whenever any one of its preferred states are matched. A second variety, *multiplicative* (AND-like) mixed selectivity, replaces the sum with a product, *M*_*ij*_ = ∏_*k*_ *f*_*ik*_(*x*_*jk*_), so the neuron activates only when all preferred states match simultaneously. Both varieties have been reported in the literature^7;11–16;36;37^.

The two cases give visually distinct intrinsic fields on the product of the two circles. Extended Data Figure 8 shows a single neuron’s tuning under both models on the (*x*_1_, *x*_2_)-torus square: under additive tuning the field resembles two circles glued together at a point (a “cross”), while under multiplicative tuning it is a single localized bump centered at the point where the two circles intersects. The additive case is what we see in Figure 2h-i, while the multiplicative case is what we observe on the grid cell torus^38;39^.

For this paper, the distinction is crucial. When the data is genuinely multiplicative, and neurons and time bins are local on the product, the disentanglement target is that product. For additive mixed selectivity, when bidirectional locality breaks, the disentanglement target is the individual covariates *X*_1_, …, *X*_*K*_, rather than their product. This is the problem we address here. The next two sections turn bidirectional locality from a diagnostic into methods that recover the covariate geometries.

### 2.4 Coherent projections separate entangled covariates

Section 2.1 established bidirectional locality as the data-intrinsic signature of a disentangled representation. We now formalize this signature by defining a scalar measure of locality on any non-negative matrix^40^ (Methods, §4.1) and then learn a non-negative projection of the data matrix that is explicitly optimized to make it bidirectionally local.

#### Formal locality

For any non-negative matrix, define the *locality* of a row as the variance of its intrinsic field about its own barycenter, the activity-weighted mean of the columns it is active on^40^ (we use a squared-*L*^1^ weighting; Methods, §4.1). It is small exactly when the row concentrates its mass on a tight cluster of columns. That is, when the row is point-like on the column space. The locality of a column is defined symmetrically. *Bidirectional locality* is the joint condition that both measures are small; every row is local on the columns and every column is local on the rows. We think of this as a formal, intrinsic signature of a disentangled representation.

#### The coherent projection

As discussed above, typical neural data matrices are rarely bidirectionally local: their rows and columns are entangled across multiple covariates. A *coherent projection* of *M* is a non-negative weight matrix 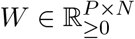 whose projected matrix

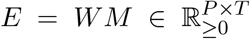

is bidirectionally local. We learn *W* by minimizing the locality loss of the rows of *E* (its modes) and of its columns (the projected time bins), evaluated on *E* itself. The rank *P* is a free parameter, chosen to be large (*P* = 100 here) so that the modes tile the targeted covariate. When *M* is driven by only one covariate, a single projection suffices. For several entangled covariates we train several projections in parallel and decorrelate them (as described below). Note that this way of optimizing for locality treats neurons and time bins asymmetrically by assuming that time bins are local within each covariate. This is true in our model of neural activity described in Section 2.2 but might not hold for other datasets. If a column does not contribute to a covariate, or contributes in a non-local way (e.g., with several “fields”), this implementation will still assign the time bin a local part of the modes.

#### Closing the loop: the cov term

Minimizing locality alone is not enough. Because locality is measured against the very matrix being optimized, the optimizer can satisfy one side cheaply without producing a genuinely bidirectionally local geometry. Specifically, on the mixed-selectivity toy data, the column locality drops below the target while the row locality stalls. The column embedding thus never recovers the latent ring (Extended Data Figure 9a,b). To rule out this shortcut we add a second, *covering* term (cov)^40^ that couples the two sides directly: where locality only asks that the columns a row loads on cluster *somewhere*, cov asks that they cluster around that row itself, and symmetrically for columns (Methods, §4.1). Optimizing loc + cov brings all four terms to target together, and the column embedding becomes a clean ring with the probe row localized on a single arc (Extended Data Figure 9c,d).

#### Implementation

The optimization we actually run adds three further ingredients, each detailed in Methods (§4.1). First, loc and cov are normalized so the target threshold is scale-free. Second, per-row and per-column violations are aggregated by a top-*k* mean, so the optimizer pushes against the worst offenders rather than the bulk. Third, and an anti-collapse floor blocks a degenerate shortcut the target alone does not.

#### Separating multiple covariates

A single coherent projection captures at most one covariate. To separate *K* entangled covariates we train *K* projections in parallel and add a decorrelation penalty that discourages any two of them from organizing the time bins in the same way, pushing each onto a different projection (Methods, §4.1.7).

#### Demonstration

We demonstrate the algorithm on the mixed selectivity toy data from Section 2.1, driven by two entangled covariates. Training *K* = 2 decorrelated projections of rank *P* = 100 and inspecting their projection matrices *E*^(*k*)^ = *W* ^(*k*)^*M*, we see that each projection recovers one covariate as a clean structure and the two projections are decorrelated onto different modes (Figure 3a–d). This works because each mode combines neurons with similar tuning to one circle while ignoring the other. If the two circles are approximately unrelated, the tuning will be spread out on the other circle, so the contribution from that circle averages out. Decoding an angle from each projection’s circle confirms the recovery quantitatively: each projection tracks one ground-truth covariate with circular correlation *ρ* ≈ 0.99 and the other near zero (Figure 3f), as the training loss settles into its target band (Figure 3e). For a demonstration on discrete states, see Extended Data Figure 10. All hyperparameters are described in Methods, §4.1.10.

**Figure 3:**
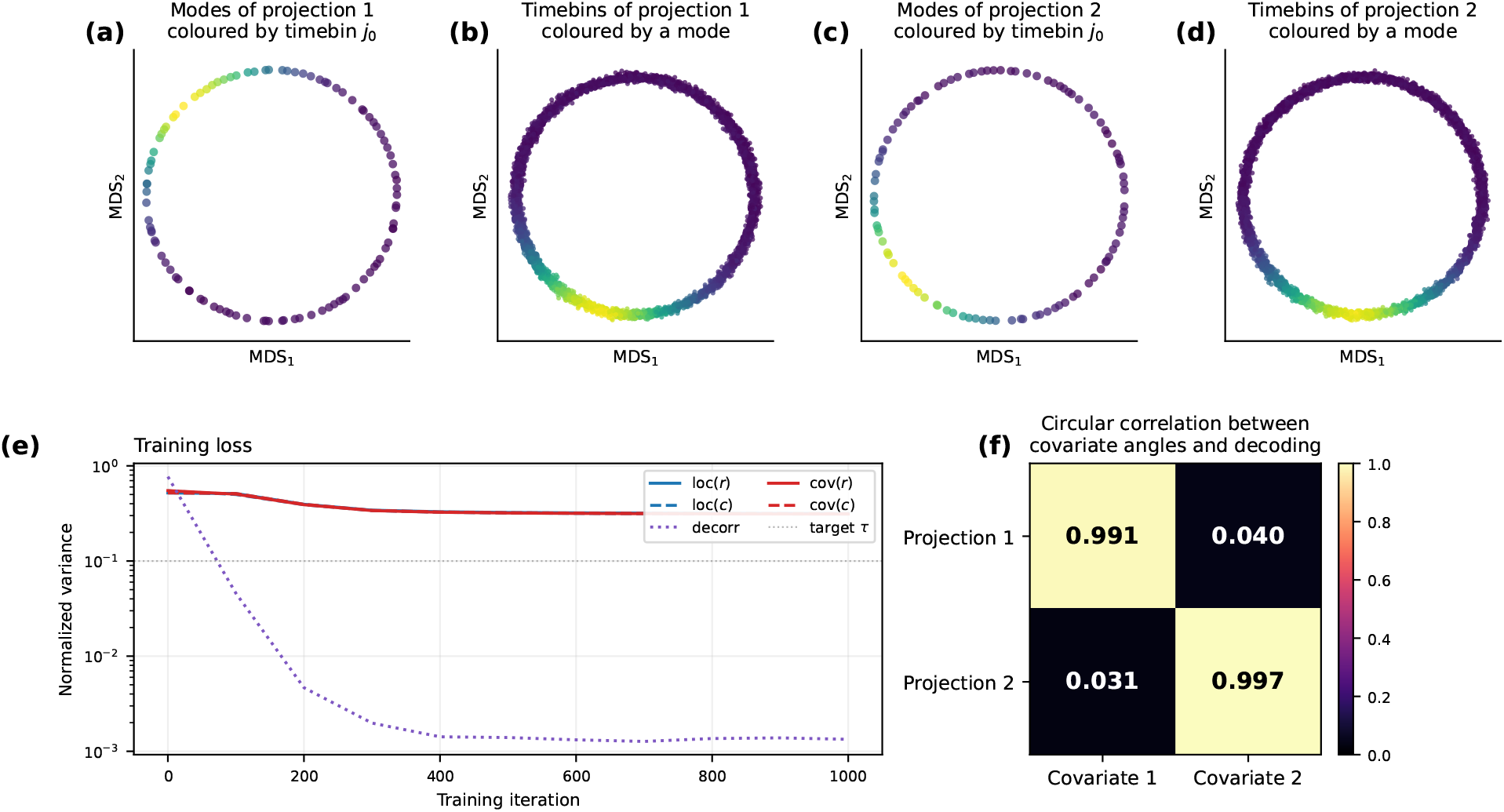
Coherent projections recover the two covariates from mixed-selectivity data. Two decorrelated coherent projections (*K* = 2, rank *P* = 100) trained on the continuous mixed-selectivity toy data of Section 2.1 (*N* = 400 all-mixed neurons, *T* = 4000, two entangled circular covariates *θ*_1_ and *θ*_2_). **(a–d)** Classical MDS embeddings (using cosine distance) of each projected matrix *E*^(*k*)^ = *W* ^(*k*)^*M* for *k* = 1, 2: its rows (the *P* modes) colored by activity at one probe timebin, *E*^(*k*)^[:, *t*_0_], (**a, c**) and its columns (time bins) colored by the activity of one probe mode, *E*^(*k*)^[*m*_0_, :] (**b, d**). Each projection recovers one circle as a clean ring. **(e)** Training loss: the locality (loc, blue) and covering (cov, red) variances of the projected rows and columns, and the cross-component decorrelation loss, driven toward the target *τ* over training (log scale). All terms converge together. In fact, all loss curves lie on top of each other. **(f)** Quantitative recovery: the projection × covariate circular-correlation matrix between a circular coordinate decoded from the time bins of each projected matrix using persistent cohomology (Methods) and the two ground-truth covariates, is near-diagonal. That is, each component tracks a different latent (*ρ* ≥ 0.99 on its own circle, ≤ 0.04 on the other). The discrete data version is in Extended Data Figure 10.

### 2.5 Clumps find small local regions on covariates

Given a matrix *M*, we can pick a set of rows **x** and columns **y** whose rectangular submatrix *M* [**x, y**] is dense, with almost every entry well above the average value in *M*. We call such a submatrix a *clump* (in reference to Good^26^). Here, we argue that a clump is local on a single covariate. That is, there is one covariate *k*^⋆^ where all time bins **y** in the clump are close and all neurons **x** in the clump have similar tuning. On the remaining covariates, the neurons and timepoints are spread out.

#### Clumps are local

While neither neurons nor time bins are point-like under additive mixed selectivity, clumps are point-like in the sense that all of the neurons **x** and time bins **y** in a clump must be close on at least one covariate. This is because an entry *M*_*ij*_ = ∑ _*k*_ *f*_*ik*_(*x*_*jk*_) is large only when neuron *i* is tuned to the state of time bin *j* in at least one covariate. A clump demands this of every row-column pair in the rectangle **x** × **y**. There are two ways to satisfy this. The straightforward way is that all rows and all columns are close on the same covariate *k*^⋆^. The rows and columns can then be tuned to anywhere on the other covariates. The alternative is that different row-column pairs rely on different covariates. This means that all the rows must be tuned similarly to more than one covariate. The probability of this coincidence decreases multiplicatively with the number of neurons and the number of covariates, if we assume at least approximate independence between the covariates. Dense rectangles in *M* should therefore be local on one covariate, motivating our search for them. A formal version of this argument for the discrete additive model is found in Methods, §4.2.1.

#### Mining clumps

Enumerating all dense submatrices is hopeless, so we mine them with an alternating fixed–point iteration^26;41^. Given an initial set of columns, we keep the rows whose standardized activity is largest on those columns. This gives a set of rows, which we use to do the symmetric update of the columns. We iterate this to a fixed point, giving us one clump. Next, we repeat this process, finding more clumps by rerunning from fresh seeds (initial sets of columns) before attempting to infer the geometry of the covariates. Because clumps find small local regions, with no global constraints, it is often challenging to sufficiently cover covariates with clumps. To solve this and ensure that we find distinct clumps, we add a soft *repulsion* away from clumps already found, pushing the iteration away from parts of *M* covered by other clumps (*K*_*c*_ = 1000 clumps used in the current examples). Repulsion is essential in the continuous case. Without it, the iteration repeatedly converges to a few of the densest regions and never covers the rest of the covariate space. After having mined many clumps, we can cluster them (here, we use DBSCAN) based on their containment overlap, and hope that the clusters correspond to the different covariates (or parts of the covariates). The update equations, standardization, repulsion penalty, and hyperparameters are in Methods, §4.2.

#### Demonstration

On the mixed selectivity toy data, each recovered clump is indeed local on one covariate: its neurons and time bins concentrate on a small region of one circle while spreading across the full range of the other (Figure 4b). To compare the clumps, we compute the containment overlap of their submatrices. This means that two clumps are close if they share many rows and columns. We first cluster the clumps (Figure 4c) and then compute an MDS embedding from the containment overlap between all clustered clumps (Figure 4d-h). Treating the clumps, rather than the rows or columns, as the point-like objects, we recover the geometry of the covariates.

**Figure 4:**
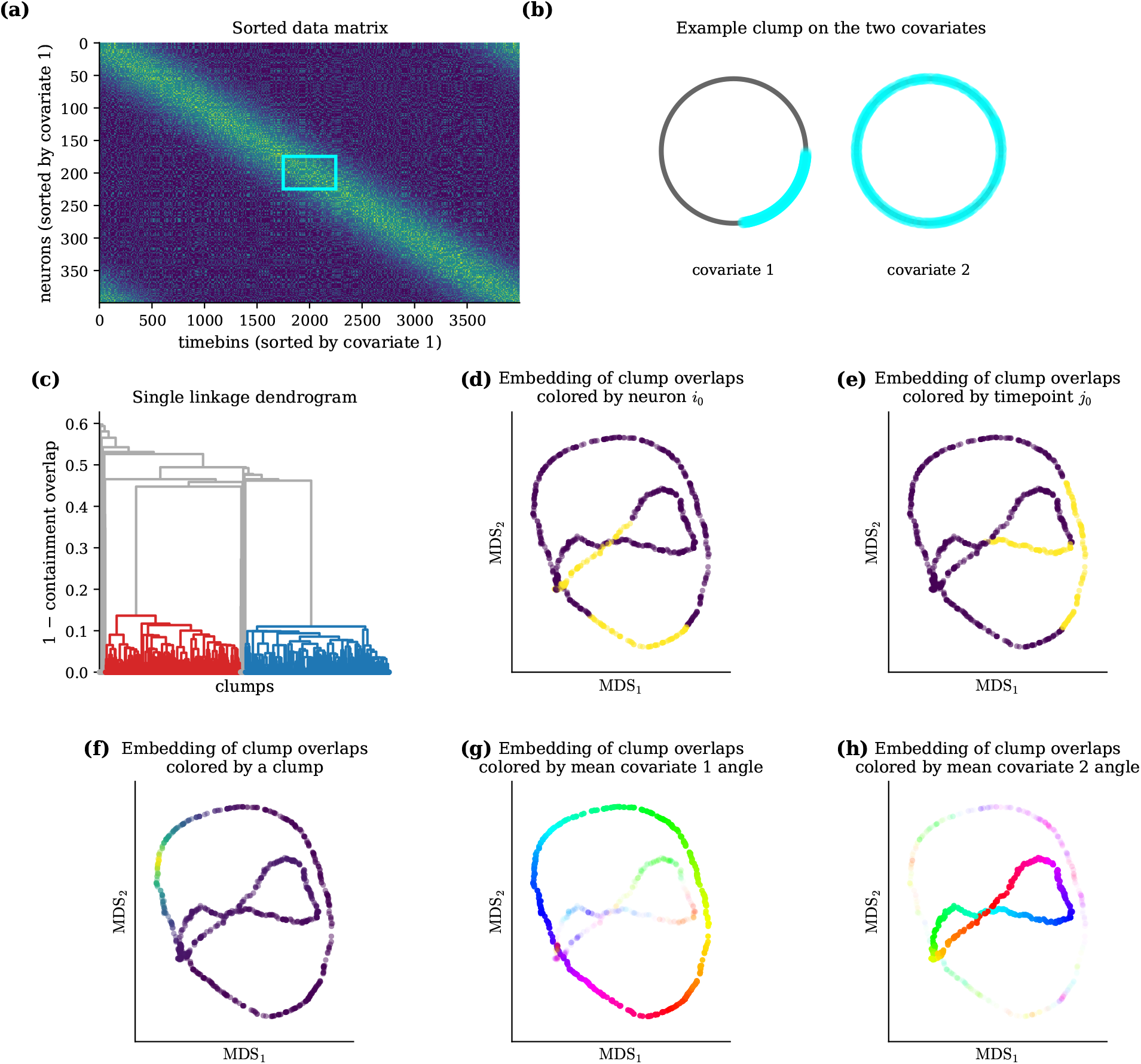
Clumps are point-like proxies for local regions on covariates. Clumps found in the continuous mixed-selectivity toy data of Section 2.1 (*N* = 400 all-mixed neurons, *T* = 4000, two entangled circular covariates *θ*_1_ and *θ*_2_). **(a)** The data matrix with rows and columns sorted by covariate 1 exposes the clumps as dense submatrices or rectangles in the matrix (cyan box, one example clump). Note, the algorithm never sees this sorting. **(b)** The neurons and time bins (both cyan) of one probe clump drawn on the two known covariate circles. It is localized on an arc of covariate 1 and spread around covariate 2. The clump has thus found all the neurons (and timebins) sharing a field in one local region of covariate 1. However, all of these neurons also have another, unrelated field, which spreads out uniformly on covariate 2. **(c)** The single linkage dendrogram on 1 minus the containment overlap between the clumps. This gives an idea of how the clumps cluster but does not reflect the clustering algorithm we use. We use DBSCAN, which we have found to be reliable. It gives the two clusters colored red and blue in the dendrogram. **(f)** The MDS embedding of clumps, colored by one probe clump. It is local on the other clumps, as it should be. **(g-h)** The MDS embedding of clumps, colored by the circular average of covariate 1 (g) or 2 (h) angles of all time bins included in a clump (i.e., the time bins indexed by **y**). Additionally, the transparency of each clump is scaled by the mean vector length of those same angles, obscuring the clumps that include timepoints that are highly variable on the covariate (because they are local on the other covariate). The discrete-data version is in Extended Data Figure 10.

#### Interpreting clumps

To relate the clumps back to the data, we can ask which clumps contain a given row or column of *M*. Coloring the containment embedding by the clump membership of a representative neuron and a representative time bin, their intrinsic field spreads over both circles (Figure 4d-e), which is exactly how we constructed the data. Next, we confirm that a probe clump is indeed local on the space of the other clumps (Figure 4f), which should mirror the covariate space. Finally, we confirm that the identified circles match the circles generating the data (Figure 4g-h). On a real dataset, relating the clumps to recorded variables in this way would be a big part of interpreting the clumps. An analogous example for discrete toy data is displayed in Extended Data Figure 11.

### 2.6 Coherent projections recover grid cell modules and their tori

Grid cells in the medial entorhinal cortex (MEC) come in discrete *modules*, each a subpopulation with its own intrinsic toroidal population activity^32;38;39;42^. A recording typically contains grid cells from several modules. This is similar to the situation with entangled columns from Section 2.1: each neuron has a localized firing field on its own module’s torus, but a population vector is a tuple containing a position on all the tori. In this way the columns entangle them while the rows keep a stacked block structure *M* = [*M*_1_; · · ·; *M*_*K*_]. We ask whether coherent projections can recover that structure, which neurons make up each module, from the neural activity alone. On the `rat r` day-1 recording from Gardner et al.^38^ (*N* = 483 cells, three modules), a single projection does not split all three at once but locks onto one. We therefore *peel* : extracting the dominant module, removing its neurons, and refitting on the remainder (Methods, §4.1.12). Three rounds recover the three modules cleanly and stably across seeds (weight-mass 0.96*/*0.93*/*0.85 on the target module, set-purity 1.00*/*0.92*/*0.94, near-diagonal confusion; Figure 5a, f and k, Extended Data Figure 12). A fourth round no longer isolates a module, its weight spreads across all three and the extracted set falls to chance purity.

**Figure 5:**
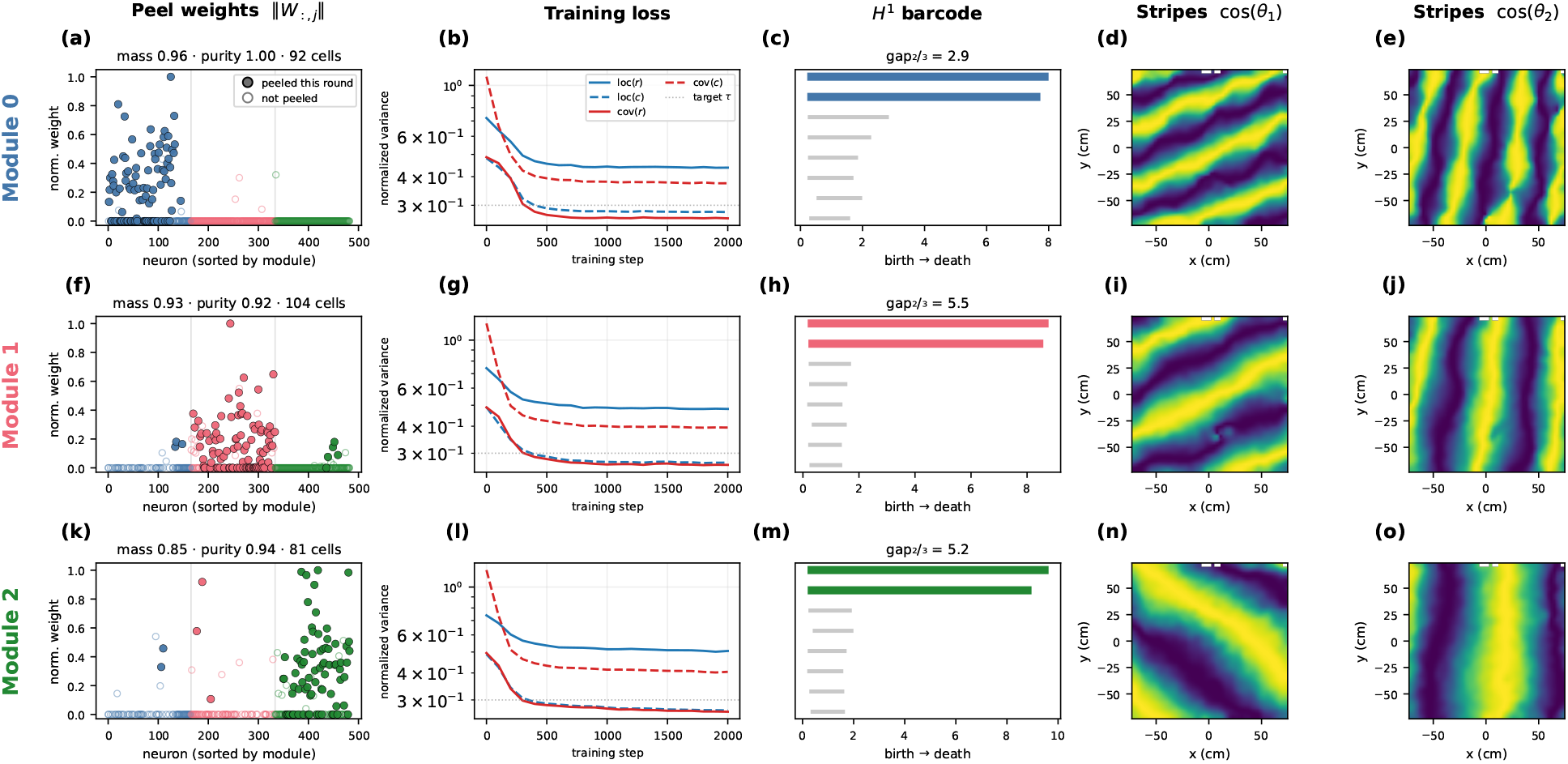
Coherent projections recover all three grid-cell modules and their tori. The grid cell modules of the `rat_r` day-1 grid-cell population of Gardner et al.^38^ (*N* = 483 neurons, three modules) is found by sequentially *peeling* off one projection or component. Each round peels one module, so row *r* is module *r* (0 → blue, 1 → red, 2 → green). Module labels are used only for coloring and scoring, never by the algorithm. Each row is one round of peeling, finding one module. All rows, panel **(a–o)**, share the same structure, described below. **Peel weights (a, f, k):** the normalized coherent weight ∥*W*_:,*j*_∥ of every still-alive neuron, ordered by held-out true module. Filled points are the neurons peeled this round and open points are kept. Each round’s component concentrates on a single module. Titles report *mass*, the fraction of the component’s total weight falling on its dominant module, and *purity*, the fraction of the peeled neurons that truly belong to it, together with the number of neurons peeled. The confusion matrix and further diagnostics are in Extended Data Figure 12. **Training loss (b, g, l):** the locality (loc) and covering (cov) variances of the projected rows (*r*) and columns (*c*) against the target *τ* (log scale). Each round converges within ~ 2000 steps. Each module’s neuron set is then formed by assigning every grid cell to the component that best explains it (Methods, §4.1.13) and analyzed with the toroidal–topology pipeline of Gardner et al.^38^. Position is used only for the speed filtering and the spatial maps. *H*_1_ **barcode (c, h, m):** persistent *H*_1_ cocycle lifetimes (death − birth). The two long colored bars are the torus’s two loops. Titles report 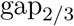, the ratio of the second-to the third-longest lifetime, which signal a torus. **Stripes (d, e; i, j; n, o):** the animal’s open-field position colored by cos of the two decoded toroidal coordinates *θ*_1_, *θ*_2_. The periodic stripes show that each module of grid cells recover a clear torus that is mapped to space. Conjunctive cells are set aside without labels (Methods, §4.1.13).

The three projections gives us the identity of the neurons from each module. If these neurons truly are a grid cell module, their joint activity should live on a torus. Running the persistent-cohomology and toroidal-decoding pipeline developed by Gardner et al.^38^ on the recovered neurons (Methods, §4.1.13), we find that every module traces a torus. That is, two long-lived *H*_1_ bars over the single *H*_2_ void expected of a torus *T*^2^. Coloring the position of the rat with the two decoded toroidal coordinates yields the expected stripe patterns with the ≈ 60° offset between the coordinates, and with the stripe period growing with the grid spacing (Figure 5). These results are reproduced in a second animal (Extended Data Figure 13). Everything is done unsupervised, from the co-activity alone. Even the (multiplicatively) conjunctive head direction and grid cells, which had to be removed from some modules, were identified without using space or head direction (Methods, §4.1.13).

### 2.7 Clumps reveal movement structure in motor cortex

Next, we show an example of how clumps can be used to find structure in datasets where we know less about what to expect. In general, this is challenging to do in an unsupervised way, partially because we must have good sampling of the covariates that drive the recorded neurons. Ideally, we would also have access to these covariates experimentally. If not, we are stuck arguing that the recovered geometry in the clumps seems non-trivial and therefore probably reflects something real. Thus, to have any hope of finding something trustworthy in a dataset, it is important that the experimental design matches the function of the recorded neurons. Here, we investigate recordings from the primary motor cortex (M1) of rats freely foraging for food crumbs in an open field^43^.

We start by finding 1000 clumps and plotting the distribution of locality scores (cov) of neurons, timebins, and clumps on all clumps (Figure 6a). We observe that, as expected, clumps have a lower locality loss, suggesting that neurons and timepoints were mixed or broadly tuned. Next, we cluster the clumps, resulting in two clusters and a group of unclustered clumps, displayed in Figure 6b. From Figure 6c and 6d, we see that most neurons and time bins are predominately assigned to neurons from one cluster. This, together with the larger locality scores, suggests that most are broadly or multi-modally tuned on one cluster. We see examples of this in Extended Data Figure 14. In Figure 6e-h we attempt to link the clumps to behavior by coloring them according to the average and standard deviation of all their time bins. The most important thing to notice here, is that cluster 2 (orange in Figure 6b) is strongly (i.e., low standard deviation) associated with very low speed and no change in head direction. Additionally, there are no noticeable changes in running or turning speed across the cluster, suggesting it operates more as a discrete state. Cluster 1 (blue in Figure 6b), in contrast, seems more interesting. We therefore study it separately in Figure 6i-l. The embeddings look like a ring where two opposing arcs are correlated with low and high speed (Figure 6i and k) and the two perpendicular opposing arcs are correlated with right and left turning. Thus, different movement patterns seem to be represented at different parts of a one-dimensional structure. Despite this, there are still parts of the cluster with a large standard deviation, suggesting that those parts might be doing something else, not directly related to running or turning speed. The identified structure expands on Mimica et al.^43^, who show how single cells are tuned to different behavioral variables. Here, we recover the organization of the neural activity without using any behavioral information. That running and turning speed shares a single continuum is not obvious, and it implies that they are related. Specifically, the rat rarely ran and turned fast at once. More generally, the ring-like structure hints that the brain often organizes neural activity into continuous structures tied to behavior, sometimes in surprising ways.

**Figure 6:**
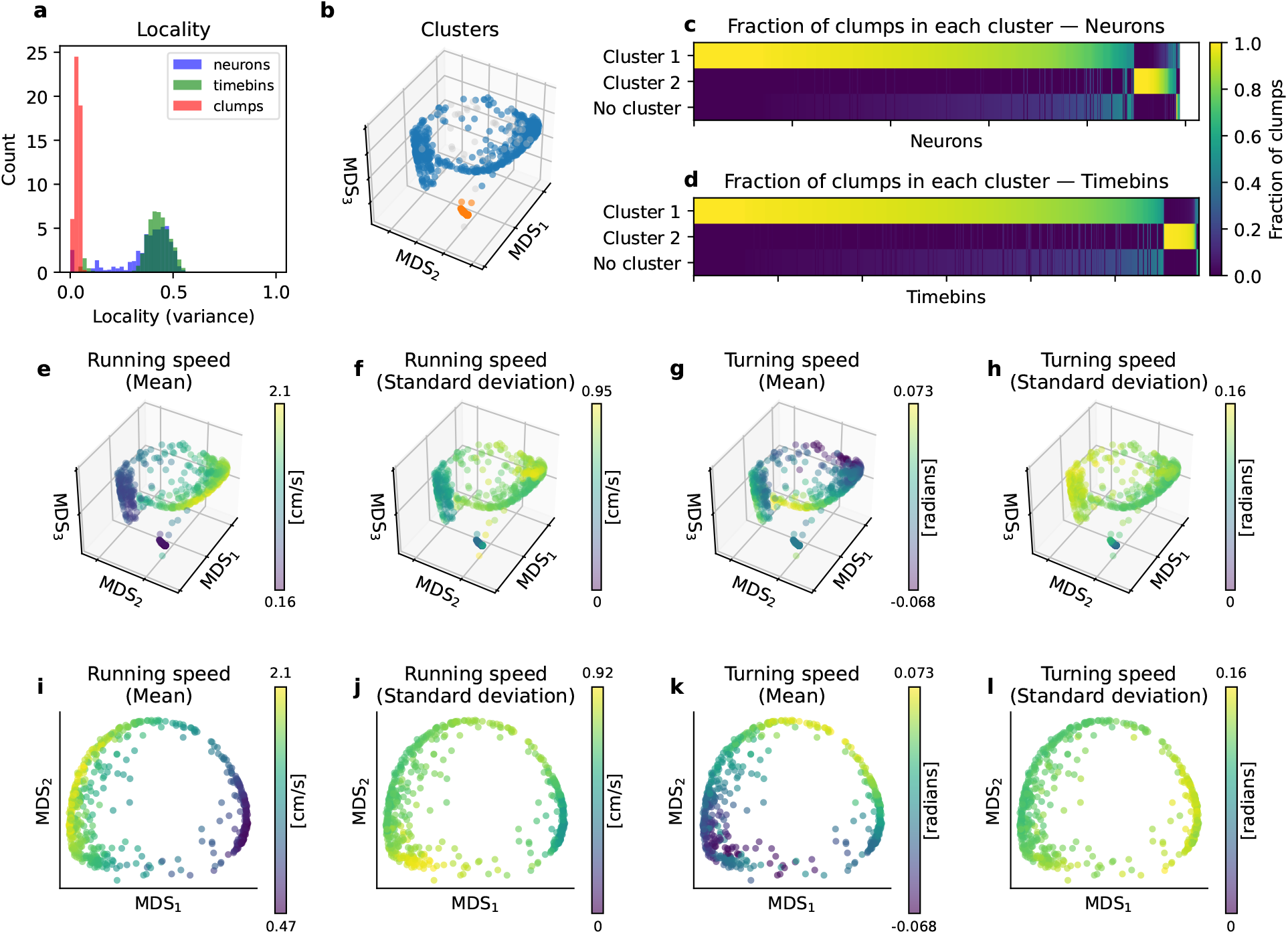
Clumps find structure in motor cortex. Clumps mined from the primary motor cortex recording of a rat foraging in an open field^43^, clustered using DBSCAN. **(a)** Distribution of locality (loc) on clumps over neurons, time bins, and clumps. For neurons and time bins, this is computed on the stacked clump matrices directly, whereas for the clumps it is computed on their containment overlap matrix. **(b)** 3D MDS embedding of all clumps from their containment overlap, colored by cluster membership. Unclustered clumps are gray. **(c–d)** Each neuron and time bin can be included in many clumps. Here we display the fraction of those clumps that belong to each cluster for every neuron (c) and time bin (d). The neurons and time bins are sorted by how strongly they belong to each cluster. **(e–h)** 3D MDS embeddings of all clumps colored by the mean (e) and standard deviation (f) of running speed, and the mean (g) and standard deviation (h) of turning speed (rate of head-azimuth change), averaged over the time bins of each clump. **(i–l)** The same four covariates on a 2D embedding of cluster 1. Running speed varies smoothly along one axis of the ring, low to high across two opposing arcs. Similarly, turning speed varies along the perpendicular axis.

**Figure 7:**
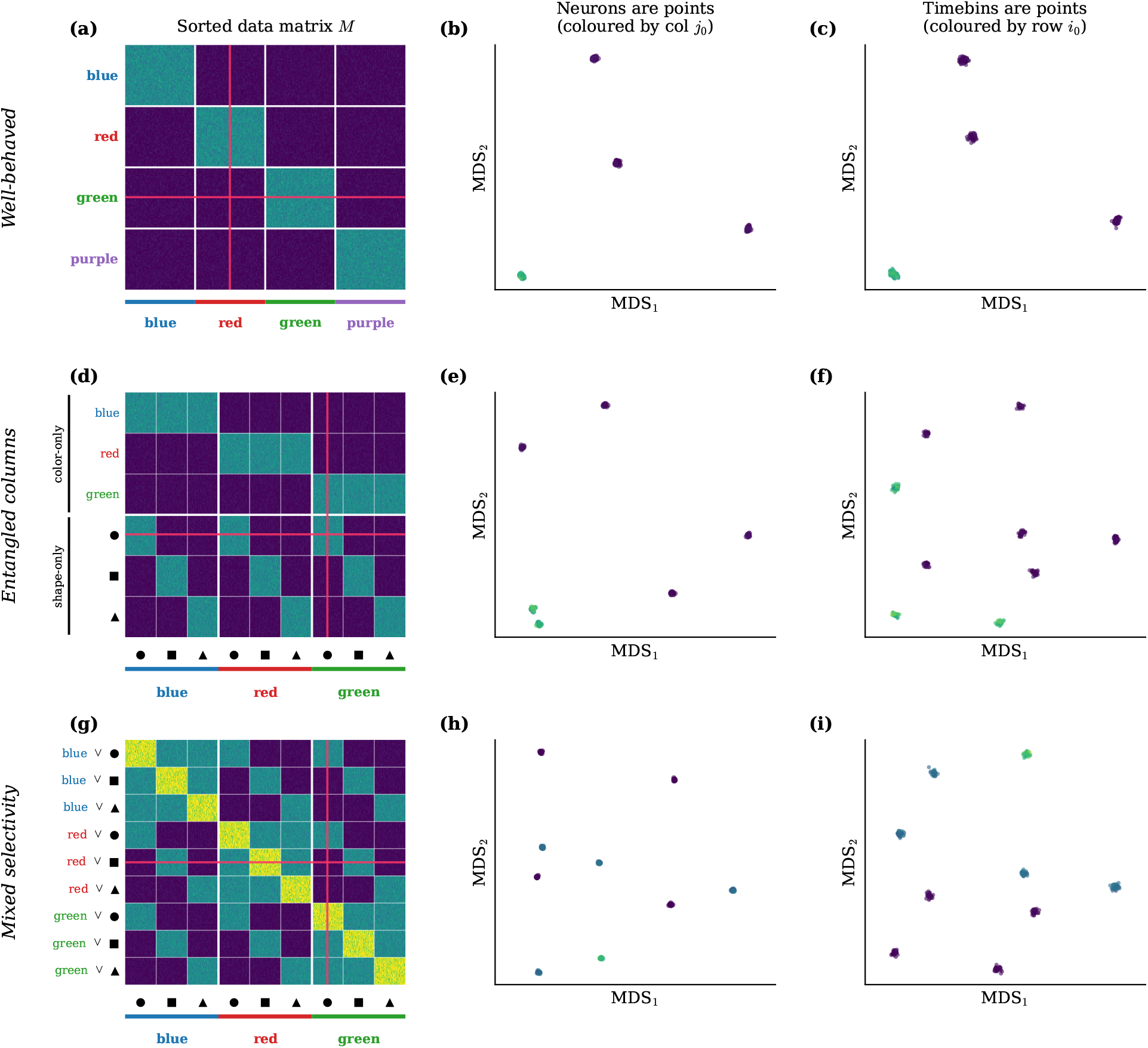
The three bidirectional-locality cases on discrete data. Discrete colours/shapes vocabulary; same 3 × 3 layout as the main-text phenomenon figure (rows: well-behaved *K* = 1, entangled columns *K* = 2 pure, mixed selectivity *K* = 2; columns: data matrix *M*, row-MDS, col-MDS). *(a–c)* one-hot tuning to four colours: four clusters in both embeddings — bidirectional locality holds. *(d–f)* colour-only + shape-only neurons: the row-MDS recovers the disjoint union (6 clusters = 3 colours + 3 shapes), the col-MDS the 3 × 3 = 9 joint states. *(g–i)* mixed (colour, shape) neurons: both embeddings show the 9-state product geometry — locality fails on both sides.

**Figure 8:**
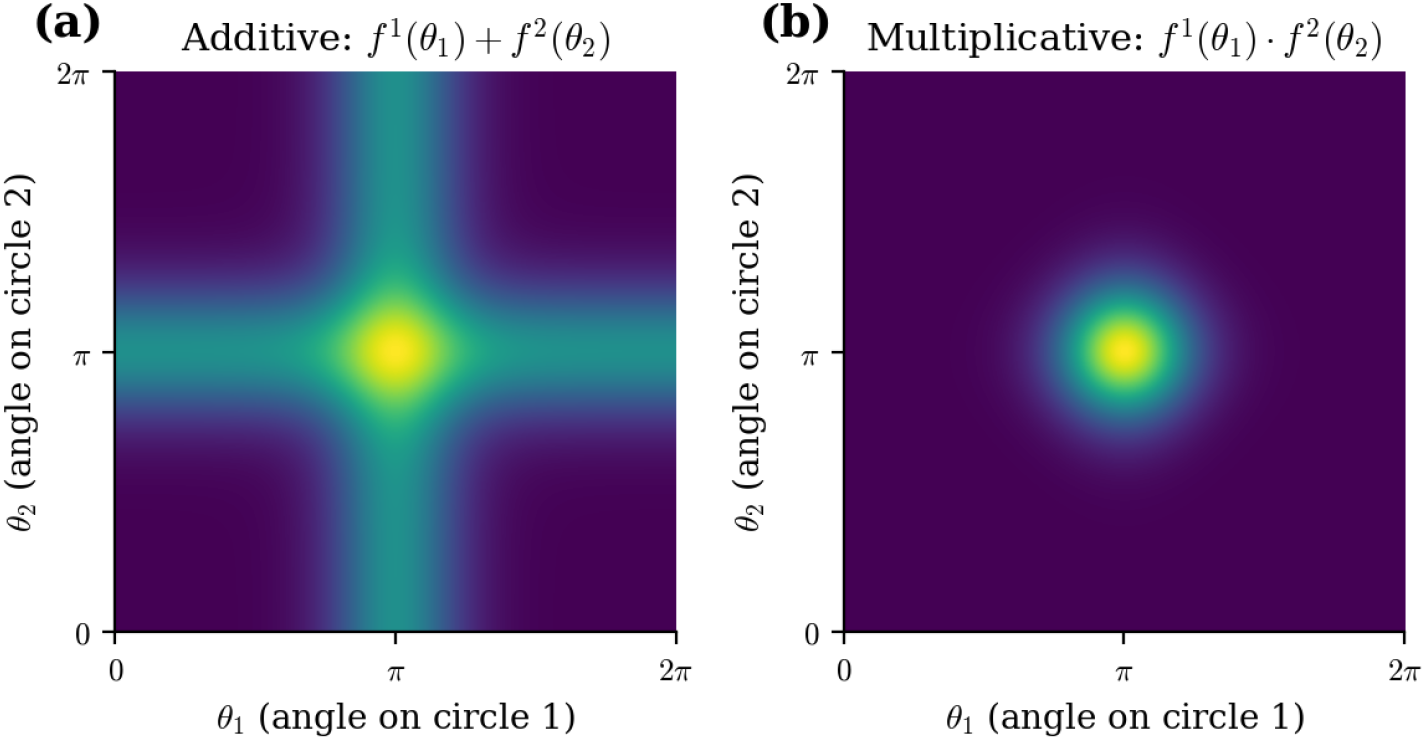
Additive vs. multiplicative mixed selectivity for one neuron tuned to two circles. Heatmaps show the firing rate of a neuron with preferred angles (*µ*_1_, *µ*_2_) = (*π, π*) as a function of the input (*θ*_1_, *θ*_2_) on the product of the two (unwrapped) circles. Tuning curves on each circle are von Mises with concentration *κ* = 4. (a) Additive, *f*_1_(*θ*_1_) + *f*_2_(*θ*_2_): each contribution lights up its own field independently, which covers the entire other circle. This gives a “cross”, which is really two circles connected at a point, the middle). (b) Multiplicative, *f*_1_(*θ*_1_) · *f*_2_(*θ*_2_): both circles must be in the field of the neuron for it to be active, so the field collapses to a single localized bump at (*µ*_1_, *µ*_2_). The methods in this paper address the additive case. Multiplicative tuning is already bidirectionally local on the product.

## 3 Discussion

Every population analysis that directly compares neurons or time bins silently assumes that they are point-like in the space we would like to recover. Disentangling the data into a couple of unrelated sources of variation is easier to interpret than looking at all of them mixed together. That is why we often find it helpful to cluster neurons or timepoints. As we have seen, this strategy only works when what is being clustered is driven by a single covariate. Bidirectional locality is a signature for whether or not the point-like assumption holds. This is the main message of the paper.

Based on this idea, we developed two unsupervised tools to get at disentanglement from different directions. Coherent projections aim to find an entire covariate at a time by learning a non-negative map that makes the projected data matrix bidirectionally local. Clumps, in contrast, find small bidirectionally local regions on the covariates by searching for dense submatrices. We showcase the potential of these methods on entangled toy data. Importantly, they manage to recover the covariates when both the rows and the columns are entangled. We applied the two methods to real neural data, revealing grid cell modules in data from MEC^38^ and a ring-like structure correlated with running and turning speed in data from M1^43^.

One important note for both of these methods is how they deal with neurons and/or time bins which disagree on how to split up the covariates. The example encountered here involves multiplicatively tuned conjunctive head direction and grid cells, firing in a grid pattern but only when facing a particular direction. The grid cell recording we used^38^ contains both pure grid cells and conjunctive ones. According to the data, it is therefore unclear if we should recover the grid cell torus and head direction circle separately or as their product (which we probably lack neurons to sample sufficiently). We believe that, in general, it is most useful to split the covariates and use the pure (or additively mixed) neurons and time bins to do so. Incorporating a way of doing this in general would be a natural next step.

Taking a step back, the problem of disentanglement is relevant in many fields, such as topic modeling^44;45^, gene expression^46;47^, and representation learning^2;48^. Furthermore, the tools we have developed here are related to many other methods, such as biclustering^46;47^, community detection^49;50^, non-negative matrix factorization^24;51^, and dimensionality reduction^21;52^. Most of them, however, only use one direction (rows or columns) at a time. These and other related ideas are discussed in more detail in Appendix 4.2.6. One particularly relevant idea is the contrast between representing covariates linearly in a euclidean space, known as the linear representation hypothesis^53;54^, and the more field-like coding we have discussed here. Bhalla et al.^5^ stress how field-like coding might be prevalent in large language models. The ideas and tools developed here build on the assumption of field-like tuning and might thus be useful for interpreting artificial neural networks in which the linear representation analysis fails. Finally, we suspect that the question of what to represent as points can clarify a wide range of problems, both inside and outside of neuroscience.

## 4 Methods

### 4.1 Coherent projections: full algorithm

In this section we detail the coherent–projection algorithm in the exact form used to produce Figures 3, 9, and 10. Section 2.4 introduced the conceptual loss loc + cov on rows and columns of the projected matrix *E* = *WM* ^40;55^. We explain all technical details of these constructions like the explicit formulas for the locality and coherence loss. We also explain several tricks that are needed in order to guarantee a well behaved optimization.

**Figure 9:**
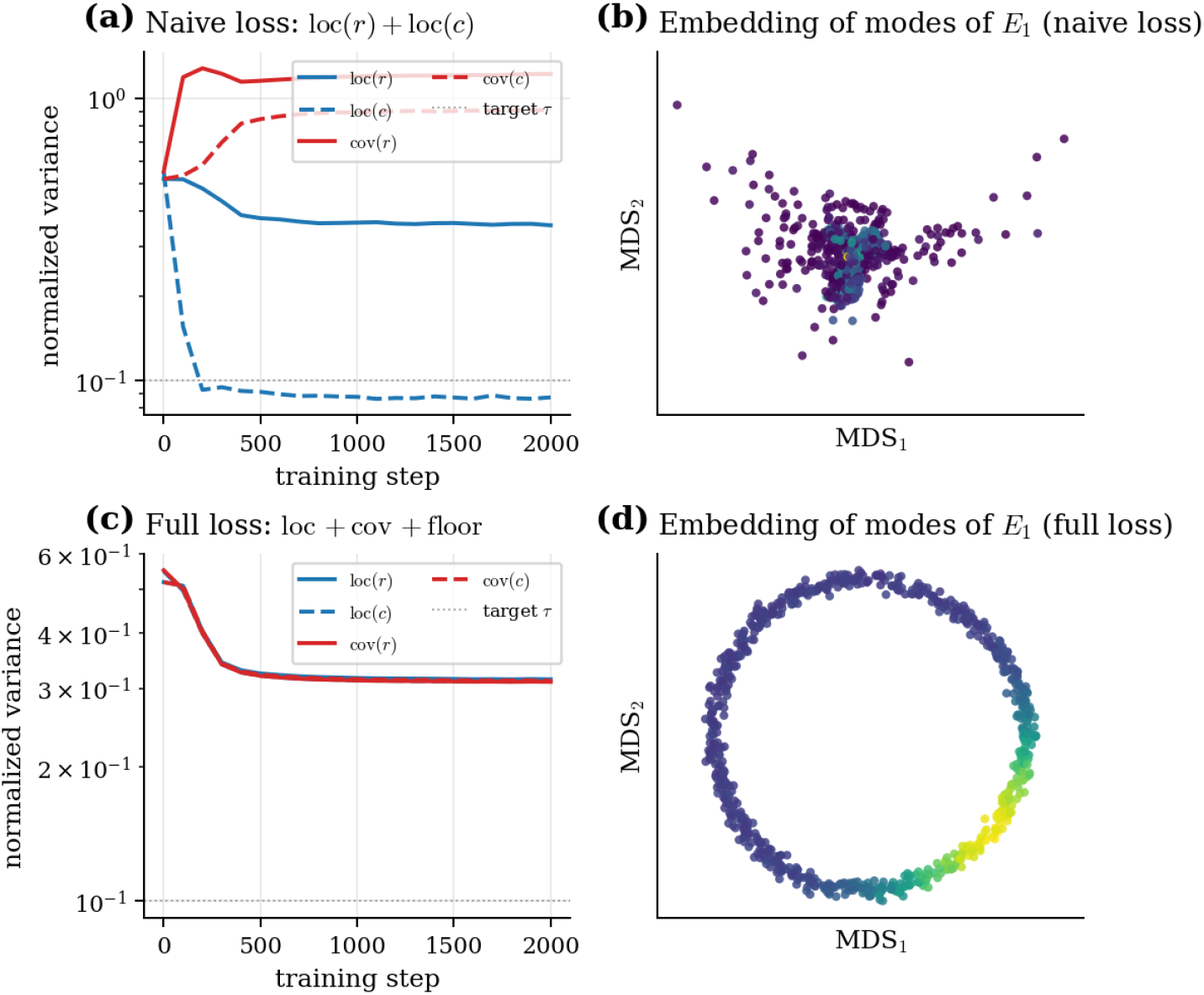
Why the covering term cov is necessary in addition to locality. Two training runs on the continuous mixed-selectivity toy data from Figure 2 (*N* = 240 all-mixed neurons, *T* = 2000, two underlying covariates). The algorithm is run with *K* = 2 decorrelated coherent projections, each of rank *P* = 100. **(a)** Naive bidirectional training, loc(*r*) +loc(*c*) only (no cov, no anti-collapse floor). The column term loc(*c*) drops below *τ* but loc(*r*) stalls and the two cov traces stay an order of magnitude above *τ*. The loss is satisfied without achieving bidirectional locality. The resulting col-MDS of the first component *E*^(1)^ = *W* ^(1)^*M*, colored by the activity of a probe mode *E*^(1)^[*m*_0_, :]. The point cloud is not a ring, so the circular covariate has not been recovered. Full training, loc + cov +floor. All four traces converge to *τ* together. **(d)** The corresponding col-MDS of *E*^(1)^ is a clean ring, and the probe mode’s activity *E*^(1)^[*m*_0_, :] localizes on a single arc. Bidirectional locality has been achieved and the covariate has been recovered.

**Figure 10:**
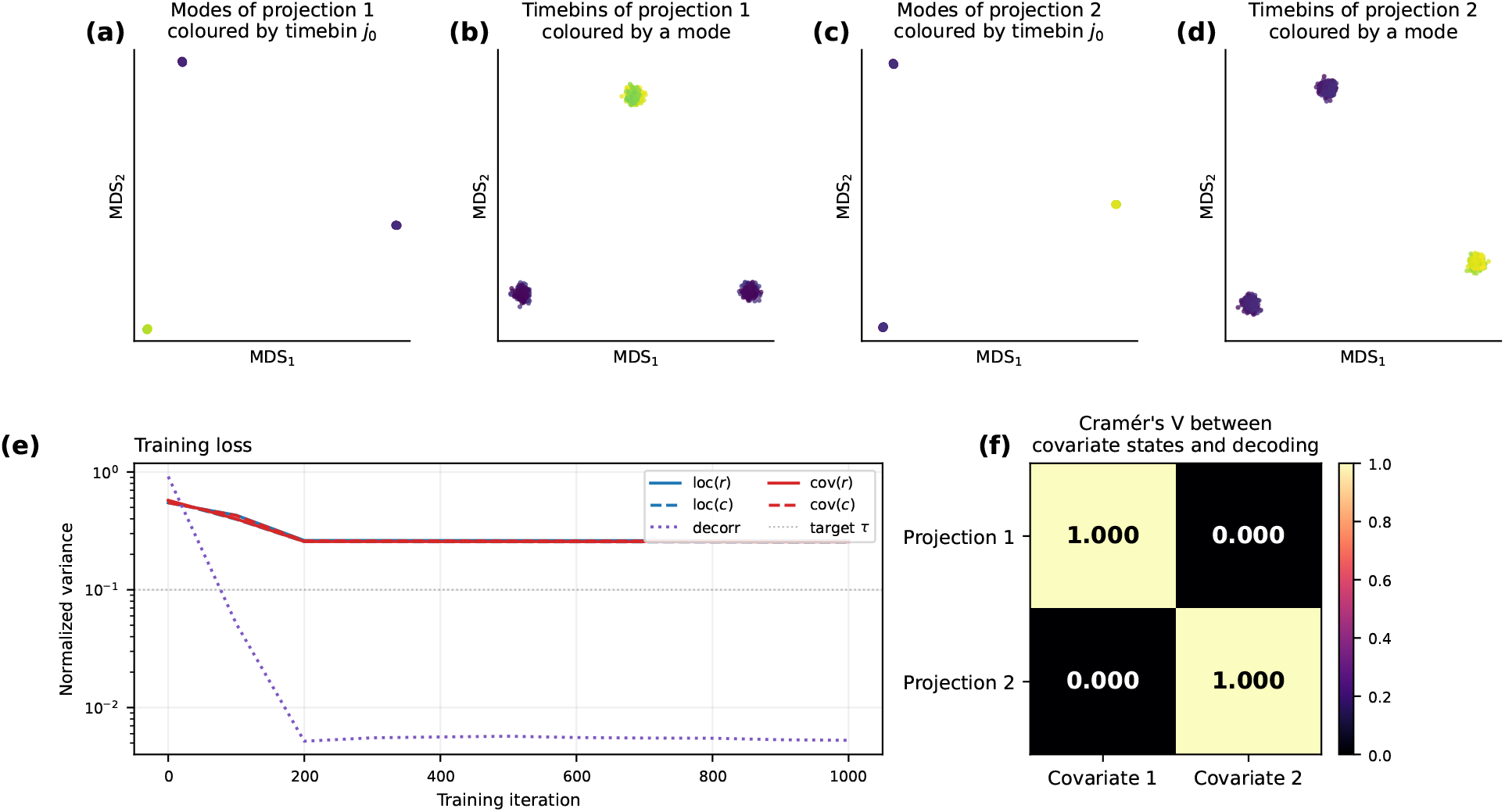
Coherent projections recover the latent factors of discrete mixed-selectivity data. The discrete-data counterpart of Figure 3: two decorrelated coherent projections (*K* = 2, rank *P* = 100) trained on the additive-mixed 3 × 3 colour/shape data of Figure 2. For each component *k*, classical MDS of the projected matrix *E*^(*k*)^ = *W* ^(*k*)^*M* — *(a,c)* its rows (the modes) and *(b,d)* its columns (timebins) — collapses to three well-separated points, the three states of one factor; the two components are decorrelated onto different factors. *(e)* Training loss: the locality and covering variances and the cross–component decorrelation converge to target together.

#### 4.1.1 Setup

Let 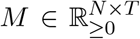 be the data matrix (we assume non–negative activities; otherwise replace *M* by max(*M*, 0) or another non–negative transform). A *coherent projection* is parameterized by a non– negative weight matrix 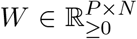, with *P* the rank of the projection (the number of *modes* per component). The projected matrix is

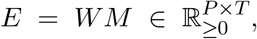

with rows *r*_*p*_ ∈ ℝ^*T*^ (the *P* modes) and columns *c*_*t*_ ∈ ℝ^*P*^ (the *T* time bins after projection). We learn *W* such that the rows of *E* are local on the columns of *E* and vice versa, in the sense made precise below.

To separate the data matrix into several decorrelated coherent projections, we instantiate *K* components *W* ^(1)^, …, *W* ^(*K*)^ in parallel and add a Mantel–style cross–component penalty (§4.1.7). Each 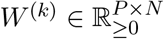 has the same dimensions; in our experiments *P* = 100 and *K* ranges from 2 to 3.

#### 4.1.2 Probability kernels via squared *L*^1^ normalization

We restate the *squared–L*^1^ kernel from Section 2.4 for the algorithm specification:

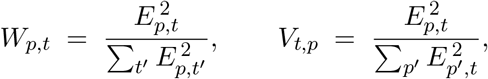

so that ∑_*t*_ *W*_*p,t*_ = 1 and ∑_*p*_ *V*_*t,p*_ = 1. The matrix *W* defines a probability distribution over columns for each row; *V* defines a probability distribution over rows for each column. Squaring before normalization makes the kernel a smooth function of *E* (the gradient is well defined at *E*_*p,t*_ = 0, in contrast to |*E*_*p,t*_|) and sharpens it: a row with a few large entries places more of its probability mass on those entries than an *L* –normalized row would, so the row barycenter 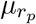 tracks the dominant peaks of the row rather than being pulled by long, broad tails.

#### 4.1.3 Row and column barycenters

With these kernels, the barycenter of row *r*_*p*_ in column space and the barycenter of column *c*_*t*_ in row space are

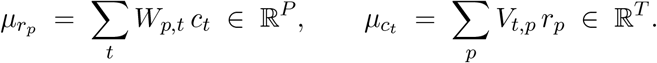

In matrix form, the stacked row barycenters are Φ = *WE*^⊤^ ∈ ℝ^*P×P*^ (the *p*–th row is 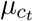) and the stacked column barycenters are Ψ = *V E* ∈ ℝ^*T ×T*^ (the *t*–th row is *µ*_*c*_). Both are computed in closed form from *E, W, V* at every optimization step.

#### 4.1.4 Locality and bidirectional locality

The four Fréchet variances that appear in the loss are

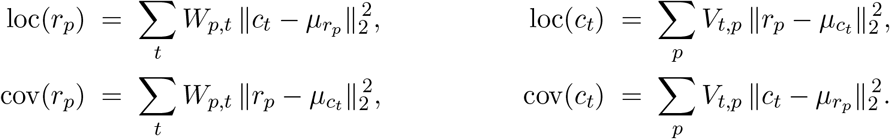

The first two (loc) measure whether the points a row (resp. column) loads on are clustered around their own weighted mean. This is the naive locality of Section 2.4. The second two (cov) measure whether the points a row (resp. column) loads on have barycenters that themselves cluster around the row (resp. column) being scored. As discussed in Section 2.4, loc alone admits a one–sided collapse, while cov closes the loop between the two sides.

The four quantities can be evaluated without forming Φ or Ψ explicitly. Expanding the square in loc(*r*_*p*_) yields

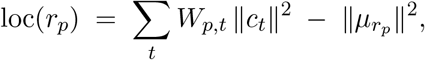

and an analogous identity for loc(*c*_*t*_), cov(*r*_*p*_), cov(*c*_*t*_). These are the closed–form expressions used in our implementation.

#### 4.1.5 Scale normalization

The variances loc and cov have units of (squared distance in row or column space), and their natural scale depends on *E*. To make the thresholds in the loss scale–free, we divide each row–space variance by the squared average pairwise distance between rows of *E*, and each column–space variance by the squared average pairwise distance between columns. Concretely, with 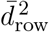 the mean squared pairwise distance between rows of *E* (estimated on a subsample of up to 256 rows at each step) and 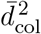 the analogous quantity for columns,

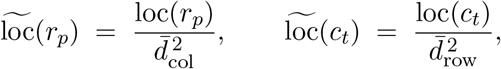

and analogously for cov. Row–space variances are normalized by the row scale; column–space variances by the column scale. A target value *τ* for the normalized variance can now be interpreted geometrically: *τ* = 0.1 means a row’s loaded columns occupy a region whose extent is roughly 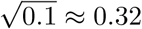 times the diameter of the whole column cloud.

#### 4.1.6 Hinge losses with a top–*k* aggregation

Given a ceiling target *τ*_ceil_ *>* 0 and a floor target *τ*_floor_ ≥ 0, we use the hinges

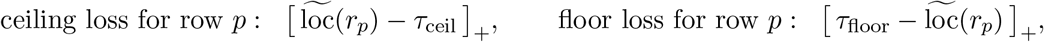

and analogously for cov and for the column terms. A naive aggregation would average these hinges over all *P* rows and all *T* columns. We instead aggregate via a *top–k mean*: we sort the hinge values across rows (resp. columns), keep the *k*_row_ largest (resp. *k*_col_ largest), and average those.

The optimizer therefore focuses its pressure on the worst–offending rows and columns at each step rather than on the bulk, which we find empirically helps the trainer escape the trivial regime where most modes are dead and a small number of modes carry all the variance. In our experiments *k*_row_ = *k*_col_ = 50.

Letting top*k*⟨·⟩ denote the top–*k* mean, the per–component variance, covering, and floor losses are

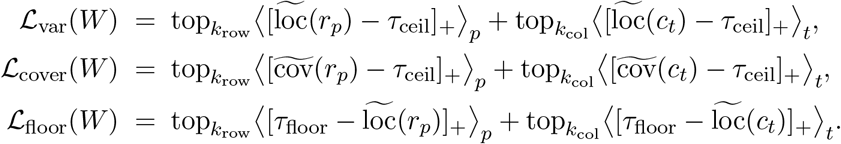

The floor is applied to the (raw, naive) 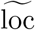 rather than 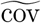: it prevents the optimizer from collapsing the column geometry of *E* to a single point or a few points, which would otherwise be a cheap way to satisfy the ceiling on cov.

#### 4.1.7 Cross-component decorrelation

When we learn *K* ≥ 2 components *W* ^(1)^, …, *W* ^(*K*)^ in parallel, we want them to capture *different* latents. We enforce this via a Mantel–style penalty on per–component column embeddings. At each step we sample a batch of *B* = 256 time bin indices *B* ⊂ {1, …, *T*}, form per-component embeddings 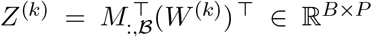 (the rows of *Z*^(*k*)^ are the projected timebins), and compute the pairwise distance matrix *D*^(*k*)^ ∈ ℝ^*B×B*^ of *Z*^(*k*)^. The mean-centered, flattened upper triangles 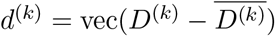 are then compared via Pearson correlation,

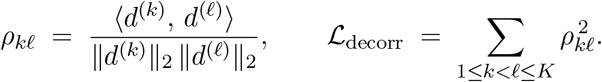

A high *ρ*_*kℓ*_ means components *k* and *ℓ* organize the same set of time bins similarly — they have caught the same latent. Squaring penalizes both same–sign and opposite–sign agreement.

#### 4.1.8 Total loss and parameterization

The total loss summed over components is

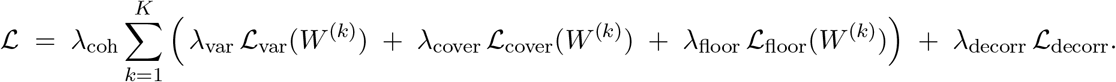

Non–negativity of *W* ^(*k*)^ is enforced both at parameter level (a ReLU is applied to the raw parameter before any use, so the effective *W* ^(*k*)^ is non–negative for free) and by an explicit projection step after every Adam update (clamping the raw parameter at 0). Each *W* ^(*k*)^ is initialized as | 0.01 · *N*(0, *I*) |.

#### 4.1.9 Optimization

At each step we sample *B* = 256 time bin indices *B*, form the batch 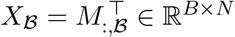, compute 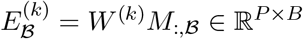 for each component, evaluate 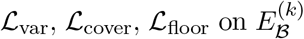, evaluate *L*_decorr_ on the batch–level embeddings, and take an Adam step on the raw parameters (learning rate 10^−3^, no weight decay) followed by a non–negativity clamp.

#### 4.1.10 Hyperparameters used in the paper

The coherent–projection runs in this paper use the following settings.

| Symbol | Meaning | Diagnostic<br>Fig. 9 | Continuous toy data<br>Fig. 3 | Discrete toy data<br>Fig. 10 | Module peel<br>Figs. 5, 13 |
| --- | --- | --- | --- | --- | --- |
| $K$ | number of components | 2 | 2 | 2 | 1 |
| $P$ | modes per component (rank of $W^{(k)}$) | 100 | 100 | 100 | 20 |
| $B$ | batch size (timebins per step) | 256 | 256 | 256 | 200 |
| steps | optimization steps | 2000 | 1000 | 1000 | 2000 |
| lr | Adam learning rate | | | $10^{-3}$ | |
| $\tau_{\text{ceil}}$ | ceiling target on normalized variance | 0.1 | 0.1 | 0.1 | 0.3 |
| $\tau_{\text{floor}}$ | floor target on normalized variance | 0.1 | 0.1 | 0.1 | 0.2 |
| $\lambda_{\text{var}}, \lambda_{\text{cover}}, \lambda_{\text{floor}}$ | weights on the three COH terms | | | 1, 1, 1 | |
| $\lambda_{\text{coh}}$ | overall scale for the COH terms | | | 1 | |
| $\lambda_{\text{decorr}}$ | weight on the Mantel penalty | 1 | 1 | 1 | 0 |
| $k_{\text{row}}, k_{\text{col}}$ | top-$k$ aggregation sizes | | | 50, 50 | |
| $\tau_{\text{emb}}$ | embedding-membership peel threshold | — | — | — | 0.2 |

#### 4.1.11 Reading the per–component embeddings

After training, each component yields a projected matrix 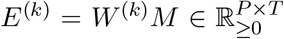. The figures in the main text display two embeddings per component:

- A *row embedding* of *E*^(*k*)^ (the *P* modes), obtained by classical MDS with cosine distance after discarding dead modes (rows with ∥ · ∥_2_ *<* 10^−9^).
- A *column embedding* of *E*^(*k*)^ (the timebins), obtained the same way, optionally on a uniform subsample of time bins to keep the eigendecomposition tractable.

Dead modes and zero columns are dropped before MDS so they do not collapse the embedding to the origin.

#### 4.1.12 Sequential module peeling

We specify the procedure behind Figure 5, in addition to Extended Data Figure 12 and 13. The idea is to use a single coherent projection to find one module at a time. That is, sequentially extract the most coherent module, remove the neurons that belong to it, and refit on the residual until no coherent module remains. This trick only works when neurons belong to a single covariate, which they do here.

##### Data matrix

We use the `rat r` day-1 recording from Gardner et al.^38^, which contains spike trains of *N* = 483 grid cells whose module identity (0, 1, 2; 166*/*168*/*149 cells) was determined in the original study and is withheld here except for scoring. We build the data matrix 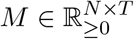 by binning spikes at 10 ms, smoothing the neurons with a Gaussian kernel (*σ* = 50 ms), applying a 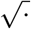 variance-stabilizing transform, and decimating the time axis by a factor of 100 (to ≈ 1 s effective resolution). We do not normalize across neurons because the locality objective is already scale-free (§4.1.5). Additionally, normalizing seemed to make it harder for first projection to lock onto a single module.

##### Extracting one module

On the current (alive) set of neurons we fit one coherent projection with a single component (*K* = 1, so *λ*_decorr_ = 0) of rank *P* = 20, using the loss and optimizer described above with target band [*τ*_floor_, *τ*_ceil_] = [0.2, 0.3], 2000 steps, and batch size 200. We restart from three seeds and keep the most coherent fit (lowest final row variance loc(*r*)). The dominant module of a fit is the module carrying the most coherent weight, 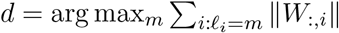, with *ℓ*_*i*_ being the held-out label of neuron *i*.

##### Peeling away the neurons from one module

Let *E* = *WM* ∈ ℝ^*P×T*^ be the projected matrix of one extracted component, which is hopefully a module. For each remaining neuron *i*, we regress its mean-centered activity *M*_*i*,:_ on the mean-centered rows of *E* and record the coefficient of determination 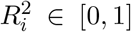, describing how well the neuron’s time course is captured by the component’s low-dimensional trajectory. Neurons with 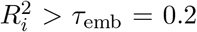 are taken as members of the extracted module and removed, while the rest pass to the next round. Peeling by embedding membership rather than by raw weight is what lets later rounds reach the weaker modules.

##### Rounds and scoring

We run one round per module, three for `rat_r` day 1 and two for `rat_q`, reading off the extracted module each round. A further round returns no additional module: the fourth round on `rat_r` day 1 spreads its weight 0.33*/*0.29*/*0.38 across the three already recovered modules and its 165–neuron extracted set sits at chance purity, 0.42 ≈ 1*/*3 (Extended Data Figure 12). For each round we report the *weight–mass* on module *m*,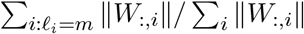 (over the alive neurons), and the *purity* of the extracted set *S*, |{*i* ∈ *S* : *ℓ*_*i*_ = *d*}|*/*|*S*|. The labels *ℓ*_*i*_ enter only these two diagnostics; the extraction and the peel never see them.

#### 4.1.13 Toroidal topology of the recovered modules

Given coherent projections, we want to check if they have the expected toroidal topology. To do this, we rely heavily on the pipeline developed by Gardner et al.^38^.

##### Toroidal decoding pipeline

This is adapted from the method developed by Gardner et al.^38^. On a module’s neurons we apply, unchanged, their toroidal-topology pipeline: restrict to moving periods (speed *>* 2.5 cm/s), keep the (1.5 × 10^4^) most active timebins, *z*-score and reduce to six principal components, density-downsample to a landmark cloud (1.2–1.5 × 10^3^ timebins), build the fuzzy-graph metric (from 800 neighbors), compute persistent cohomology (Vietoris–Rips; *H*_1_ with coefficients in F_47_, the *H*_2_ void on a ≤ 500-point subsample), and read out the two most persistent *H*_1_ cocycles as toroidal coordinates with their cohomological / population-vector decoder. A module is a torus when two *H*_1_ bars are long relative to the rest with a single *H*_2_ bar present. We summarize the former by the second-to-third *H*_1_ lifetime ratio, gap_2*/*3_. See^38^ for details. Conjunctive (head direction-modulated) grid cells are set aside where needed (for Figure 5). However, unlike in Gardner et al.^38^, we identify them in an unsupervised fashion instead of using a manual classification, as described below.

##### Finding the full set of neurons in a module

The peel (§4.1.12) find the neurons most strongly tuned to each module; its core. To get a more accurate toroidal decoding, we expand these cores. This is done by assigning every cell to the peeled component whose projected trajectory *E* = *WM* best explains its time course, the same embedding-membership *R*^2^ used in §4.1.12, now taken as an arg max over the recovered components rather than a per-round threshold. This captures the weaker and mixed cells that an individual peel round leaves behind. For `rat r` module 0 it was further necessary to remove conjunctive head direction cells in order to recover the grid cell torus (see below). The quality of the recovered tori are 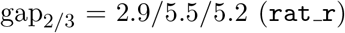 and 5.6*/*2.0 (`rat_q`). Running the same pipeline on the ground-truth pure cells gives 4.4*/*3.9*/*4.6 and 4.6*/*2.1, respectively.

##### Identifying conjunctive grid- and head-direction cells

Some grid cells are conjunctively, and multiplicatively (as in Section 8), tuned to head direction. They then fire as grid × head-direction and thus live on *T*^2^ × *S*^1^ rather than the torus *T*^2^. Keeping a few of them is harmless, as for module 1 (3%) and 2 (11%) in `rat_r`. However, there were a lot of them in module 0 (44%), disrupting the torus (even using the labeled cells). Here, we detect and remove them from the coherent projections (not the subsequent toroidal decoding) using the correlation structure between cells. For each module, we project off the top six temporal singular vectors (that the grid manifold the persistence analysis keeps) and rectify the residual, then fit one coherent projection (rank 20) on the pooled residuals, which lock onto the head direction ring. A cell is classified as conjunctive if its ring membership (*R*^2^ on the ring trajectory, as above) is large. The decoded ring angle tracks head azimuth (circular correlation 0.49) and ring membership separates conjunctive from pure cells (AUC 0.92, versus 0.68 from projection weight alone). This was only necessary on module 0 in `rat_r`. `rat_q` had almost exclusively pure grid cells, so the ring described above was not found there.

### 4.2 Clumps: full algorithm

In this section we describe the clump algorithm used to produce Figure 4, 6, and 11. A clump is a pair (**x, y**) ∈ {0, 1}^*N*^ × {0, 1}^*T*^ of binary row and column masks such that the rectangular submatrix *M* [**x, y**] is denser than the rest of *M* in a sense made precise below. We first formalize, in an elementary discrete model, why such dense blocks are local on a single covariate (§4.2.1). We then find clumps one at a time by an alternating fixed-point iteration, while enforcing diversity across clumps with a soft repulsion against previously found clumps (§4.2.2), before clustering the resulting set of clumps by containment overlap (§4.2.3), and embedding those clusters with classical MDS (§4.2.4).

**Figure 11:**
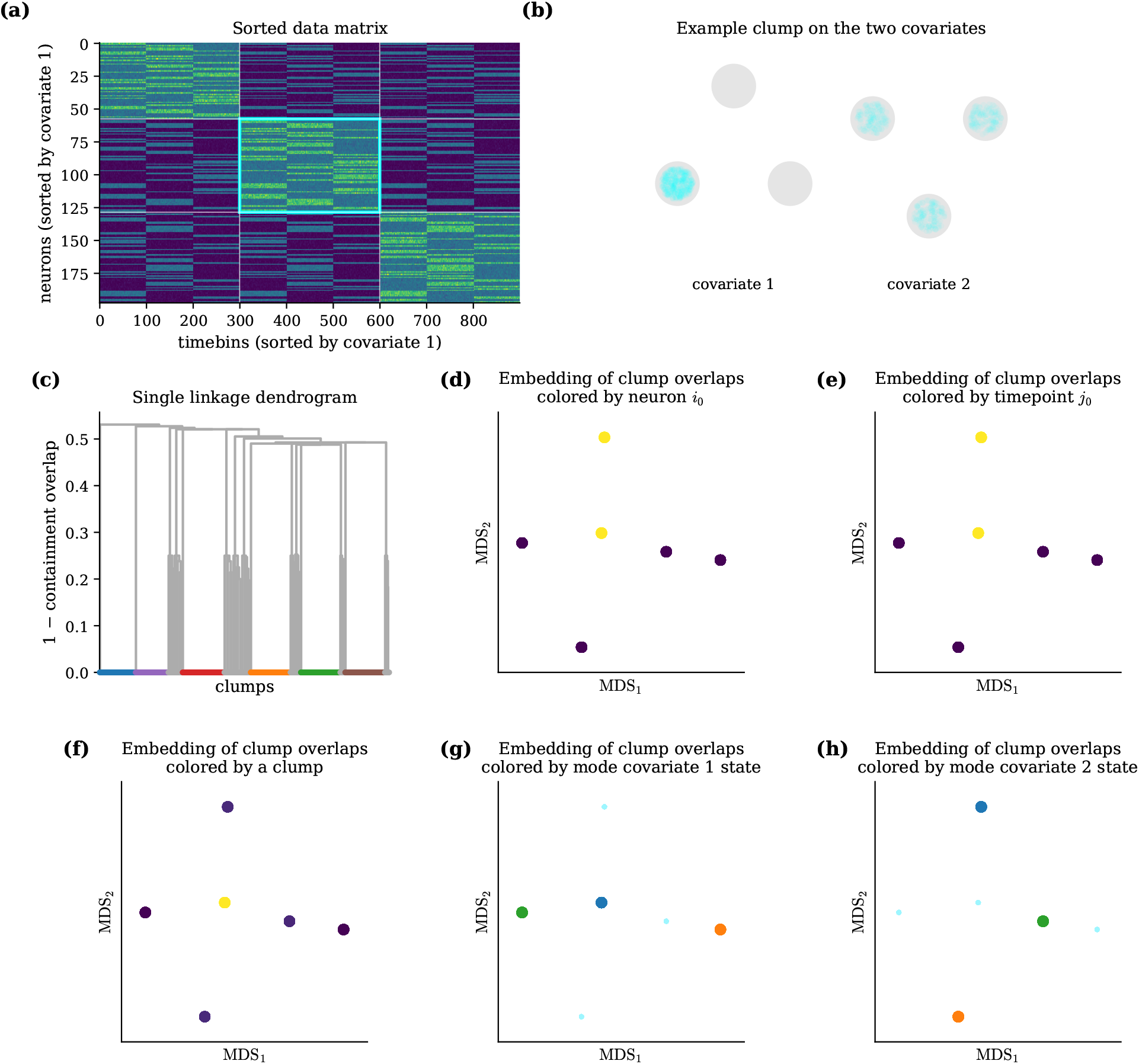
Clumps on discrete mixed-selectivity data. The discrete-data counterpart of Figure 4 (additive-mixed 3 × 3 color/shape data). Panels as in Figure 4: **(a)** the matrix sorted by covariate 1 (dense diagonal blocks, cyan box on one clump); **(b)** one example clump on the two known latents, localized at one state of covariate 1 (b1) and spread across all three states of covariate 2 (b2); **(c–e)** row, column, and containment MDS of the clump bank, each recovering six clusters (three states × two covariates); **(f)** the containment embedding colored by one example neuron (f1) and one example time bin (f2).

**Figure 12:**
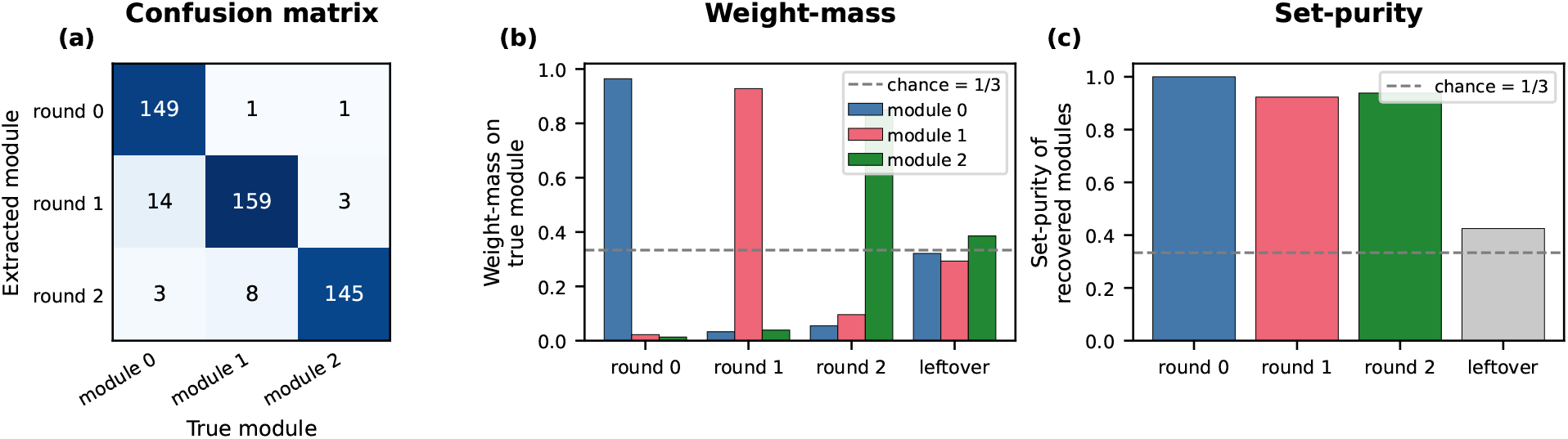
Peeling diagnostics for the rat_r day-1 modules (companion to Figure 5a). Scored against the held–out Gardner et al.^38^ module labels. Each round of the peel extracts one module and a fourth round was run past the three modules to expose the stopping signal. **(a)** Extracted-versus-true confusion matrix for the three modules. The near-diagonal structure shows each round captures a distinct module. **(b)** Component weight-mass on each true module, per round. The three rounds sit near 1 on a single module (dashed line is chance level 1*/*3), while the leftover round spreads its weight across all three (0.33*/*0.29*/*0.38). **(c)** Set-purity of each extracted module; the fraction of the extracted cells that truly belong to the round’s dominant module. The three module rounds sit far above chance (1*/*3, dashed) and the leftover round falls close to it (0.42 ≈ 1*/*3).

**Figure 13:**
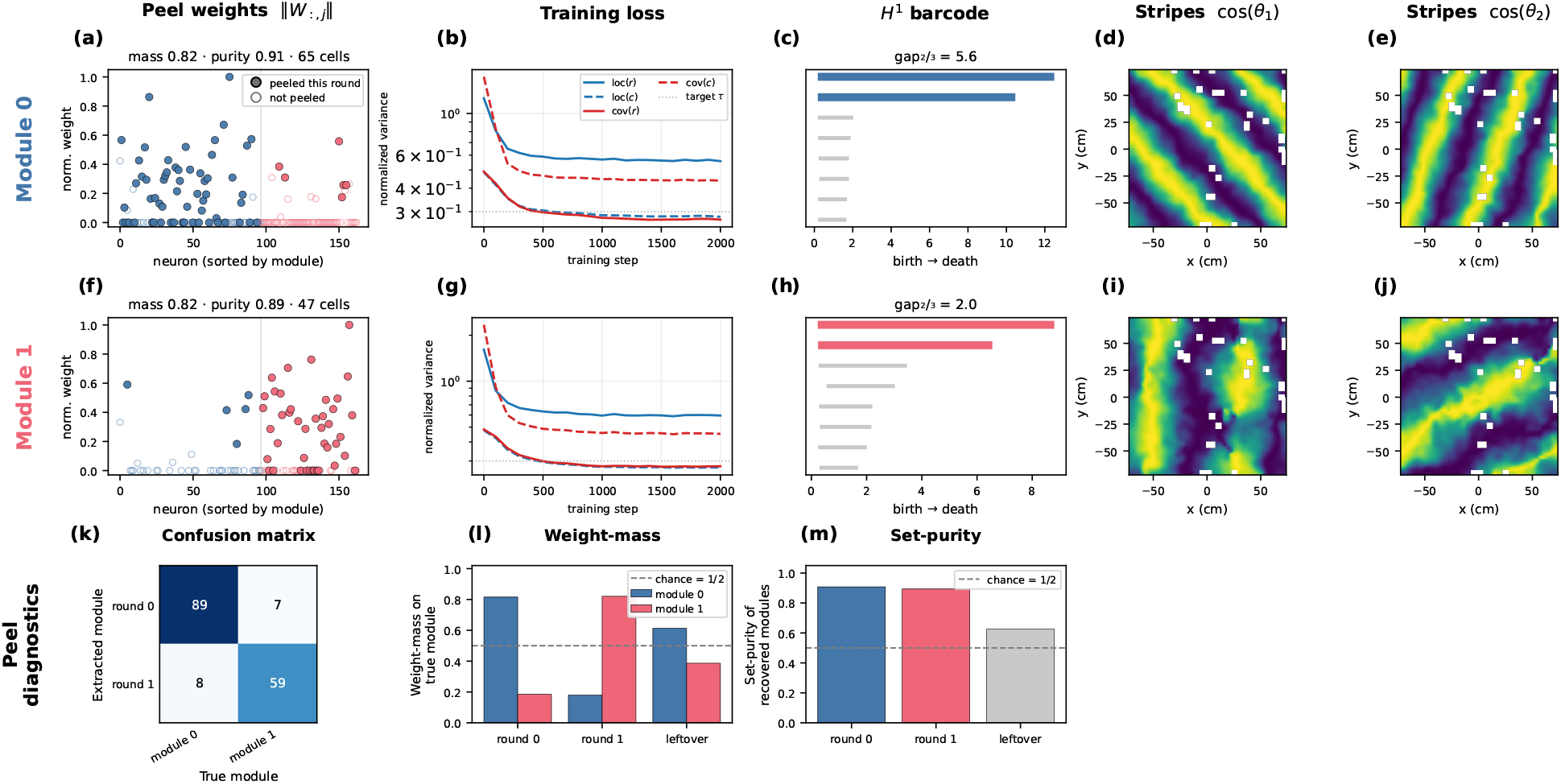
Coherent projections recover rat_q grid-cell modules and their tori. The rat q grid–cell population of Gardner et al.^38^ (*N* = 163 neurons, two modules), analyzed exactly as in Figure 5 and 12. The analysis is restricted to the open field session. **(a, f)** Peel weights: the normalized coherent weight ∥*W*_:,*j*_∥ of every still-alive neuron, ordered by held-out true module. Filled points are peeled this round, open points kept. Titles report mass (fraction of the component’s weight on its dominant module), purity (fraction of the peeled neurons truly in it), and the neuron count. **(b, g)** Training loss: the locality (loc) and covering (cov) variances of the projected rows and columns (*c*) against the target *τ* (log scale), converging within ~2000 steps. **(c, h)** *H*^1^ barcode: persistent *H*^1^ cocycle lifetimes. Titles report 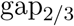. **(d, e; i, j)** Stripes: the animal’s open-field position colored by cos of the two decoded toroidal coordinates *θ*_1_, *θ*_2_. The periodic stripes show each module’s torus mapped to space. **(k)** Extracted-versus-true confusion matrix for the two module rounds. **(l)** Component weight-mass on each true module per round, with the 1*/*2 chance line. The last round spreads across both modules (61*/*39). **(m)** Set-purity per round is far above chance (1*/*2, dashed) for the two module rounds and at chance for the last round. `rat_q` is almost entirely pure grid cells, so no conjunctive cells are set aside (the head-direction ring of Methods §4.1.13 is not found here).

**Figure 14:**
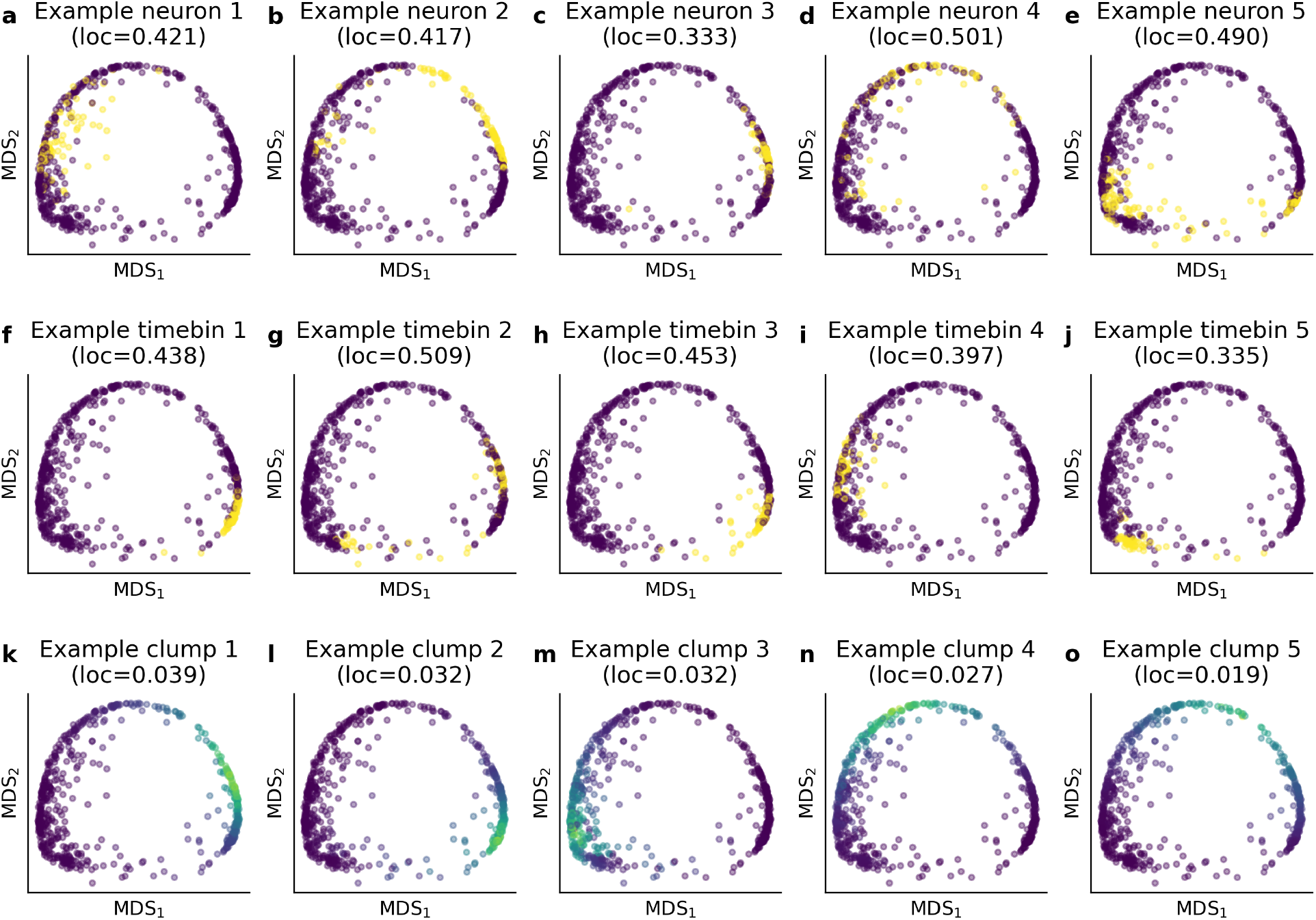
Intrinsic fields of example neurons, timebins, and clumps on the motor-cortex cluster described in Figure 6. Each panel shows the 2-D containment-overlap MDS embedding of one clump cluster (cluster 1 of Figure 6i–l), colored by the intrinsic field of a single probe. (a–e) Each clump is colored by whether it contains a neuron (i.e. the neuron’s membership across the clumps). (f–j) time bins similarly colored by clump membership. (k–o) Clumps colored by their containment overlap with the probe clump. Panel titles give each probe’s locality (intrinsic-field variance) over all clumps.

#### 4.2.1 Why dense blocks are local on one covariate

This subsection formalizes the heuristic of Section 2.5: a dense sub–block of *M* is, with overwhelming probability, local on a single covariate. We present the argument in two steps. First, every agreement between a row and a column is due to a shared state *s* on a covariate *k*, giving *KS* covariate-state bins. A dense block has so many agreements that one of the *KS* covariate-state bins must account for a fixed fraction of the block, the anchor. Second, any row or column that tries to avoid this bin has to independently agree with all rows and columns in the anchor on other covariate-state bins, which is exponentially unlikely. We give the argument for the discrete additive model, the simplest setting that isolates the phenomenon, at the minimal level *θ* = 1.

##### Model

There are *K* covariates, each taking one of *S* states. Neuron *i* has a preferred state *c*_*ik*_ ∈ {1, …, *S*} on covariate *k*, and time bin *j* has state *x*_*jk*_ ∈ {1, …, *S*}. Additive (OR–like) tuning with a binary kernel *f*_*k*_(*c, x*) = 1 [*c* = *x*] makes

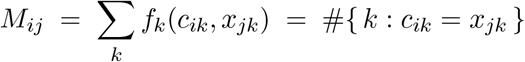

count the covariates on which neuron *i* and time bin *j* agree. Under the null in which the *c*_*ik*_ and *x*_*jk*_ are independent and uniform, each entry is Binomial(*K*, 1*/S*) with mean *K/S*. That is, a typical pair agrees on a small number of covariates by chance (i.e., without being tuned similarly), which is the sparse background against which clumps stand out.

##### Clump and locality

Fix a row set **x** (*a* = |**x**|) and a column set **y** (*b* = |**y**|). The resulting block is a clump at level *θ* if *M*_*ij*_ ≥ *θ* for every pair (*i, j*) ∈ (**x, y**). The most interesting, and most likely, level is *θ* = 1, where every pair is co-active on at least one covariate. The clump is local on covariate *k* at state *s* if *c*_*ik*_ = *s* for all *i* ∈ **x** and *x*_*jk*_ = *s* for all *j* ∈ **y**; then every pair agrees on covariate *k*, so the block is automatically a clump whatever the rows and columns do on the other covariates. (The algorithm’s mean–plus–*σ* criterion below is a soft, data–driven version of the set *M*_*ij*_ ≥ *θ*.)

##### Every clump is anchored to one covariate-state

For covariate *k* and state *s*, let *n*_*k*_(*s*) = #{*i* ∈ **x** : *c*_*ik*_ = *s*} and *m*_*k*_(*s*) = #{*j* ∈ **y** : *x*_*jk*_ = *s*}, so ∑_*s*_ *n*_*k*_(*s*) = *a* and ∑_*s*_ *m*_*k*_(*s*) = *b*. This means that *n*_*k*_(*s*) *m*_*k*_(*s*) is the number of row-column pairs in the clump that share state *s* on covariate *k*; those rows and columns are *co-local*. Summing *M*_*ij*_ over the block and exchanging the order of summation gives the exact identity

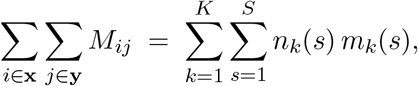

because a row and a column agree on covariate *k* exactly when both occupy a common state *s*. The clump condition *M*_*ij*_ ≥ *θ* makes the left side at least *θab*, while the right side has only *KS* terms, so

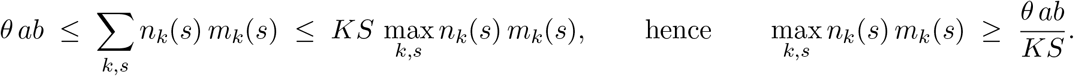

Since *n*_*k*_⋆ (*s*^⋆^) *m*_*k*_⋆ (*s*^⋆^) is the number of row-column pairs in the clump that share state *s*^⋆^ on *k*^⋆^, dividing by the total clump area *ab* gives

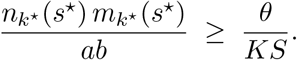

So a fixed fraction, at least *θ/*(*KS*), of the clump is *co–local*, sharing state *s*^⋆^ on covariate *k*^⋆^. Intuitively, every agreement (*M*_*ij*_ *> θ*) must be caused by one of only *KS* covariate-state bins. A dense submatrix has too many agreements (≥ *θab*) for all bins to stay light, so one bin must be heavy. This is deterministic and uses only counting. It does not yet say that the clump is entirely local: some rows and columns could still evade *s*^⋆^ and enter the clump through their other covariates. Next, we show that this is overwhelmingly unlikely.

##### Non-anchored rows and columns are improbable

Take the minimal clump level *θ* = 1, so that “matching” a column means agreeing with it on at least one covariate. The bound derived above still permits a few rows that lack state *s*^⋆^ on *k*^⋆^ yet match every column through their preferences on the other covariates. We will show that such a row is exponentially costly. Let *m*_*k*_^⋆^ = *m*_*k*_⋆ (*s*^⋆^) be the number of anchor columns; they share state *s*^⋆^ on *k*^⋆^ but are independent and uniform on the other *K* − 1 covariates. A non-anchor row *i* (with *c*_*ik*_⋆≠ *s*^⋆^) must match all *m* anchor columns to remain in the clump, and can do so only by happening to align on one of the other *K* − 1 covariates. This has a probability of 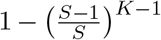 per column and is independent across all anchor columns. The non-anchor row therefore matches all *m*^⋆^ anchor columns with probability at most

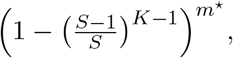

which for *K* = 2 specializes to (1*/S*)^*m*^⋆. The expected number of surviving non-anchor rows is then at most *N* (1*/S*)^*m*^⋆, which drops below one as soon as *m*^⋆^ *>* log_*s*_ *N*. The same argument applied to the transpose bounds the expected non-anchor columns by *T* (1*/S*)^*n*^⋆, below one once *n*^⋆^ *>* log_*S*_ *T*, where *n*^⋆^ = *n*_*k*_⋆ (*s*^⋆^). Past that modest size a clump has, in expectation, no non-anchor rows — and, symmetrically, no non-anchor columns — so it is exactly local on a single covariate. Letting *θ >* 1 only sharpens this: a non-anchor row would then have to agree on at least *θ* of its *K* − 1 remaining covariates per column, a rarer event. This of course becomes impossible when *θ > K* − 1. This is the precise sense in which dense submatrices are worth mining: they are proxies for local regions of one covariate.

#### 4.2.2 Mining clumps

The standardization of *M* and the fixed-point iteration (without repulsion) were introduced by Bergmann et al.^41^.

##### Dual standardization of

*M*. To make the row and column thresholding steps symmetric in scale, we precompute two standardized copies of *M* :

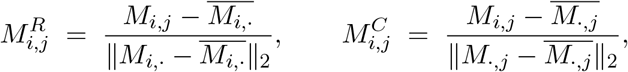

Where 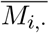, is the row mean and 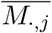 the column mean. *M* ^*R*^ has unit-norm zero-mean rows and is used when scoring rows from a column mask, while *M* ^*C*^ has unit-norm zero-mean columns and is used when scoring columns from a row mask. This double standardization decouples the row and column statistics from the absolute scale of *M* so that the thresholds *t*_*r*_, *t*_*c*_ have the same meaning regardless of how active individual neurons or time bins are.

##### Single-clump fixed-point iteration

Given thresholds *t*_*r*_, *t*_*c*_ *>* 0, a single clump is found by the iteration

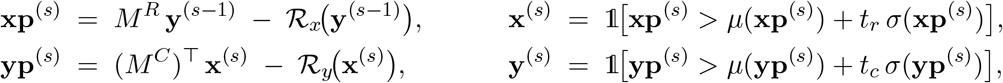

where *µ*(·) and *σ*(·) are the empirical mean and standard deviation, 1[·] is taken element-wise, and ℛ_*x*_, ℛ_*y*_ are the repulsion terms defined in introduced below (zero for the first clump). The row score 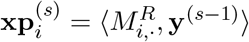 measures how strongly row *i*’s standardized activity correlates with the current column mask. Equivalently, the sum of standardized activities of row *i* on the columns selected by **y**^(*s*−1)^. The threshold *µ* + *t*_*r*_*σ* keeps only rows that are outliers above the population mean of **xp**; small *t*_*r*_ admits many rows, large *t*_*r*_ admits only the most extreme ones. The column update is symmetric.

##### Initialization

The iteration could be initialized by **x**^(0)^ or **y**^(0)^. We use **y**^(0)^, sampled i.i.d. Bernoulli(0.5) over {0, 1}^*T*^.

##### Convergence

Iteration continues until both score vectors **xp** and **yp** stabilize according to some tolerance *τ*_conv_. Letting *d*_cos_ denote cosine distance, the stopping criterion is

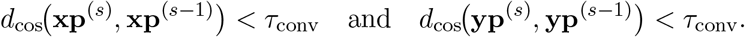

If at any step 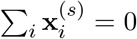 or 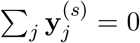 (the threshold killed the whole population), or if the iteration reaches *S*_max_ steps without satisfying the convergence test, the run is restarted from a fresh random **y**^(0)^. The same repulsion state is passed in, so a restart is just a new initialization. In our experience, only about 2–10 iterations are needed for convergence, unless a re-initialization is needed.

##### Repulsion against previously found clumps

When mining the *k*-th clump, the score updates are penalized by overlap with all *k* − 1 previously found clumps. The idea is to discourage clumps from picking the same row-column pairs, which helps cover continuous covariates with overlapping clumps. Let **R** ∈ {0, 1}^*N×*(*k*−1)^ stack the row masks of the existing clumps as columns and **C** ∈ {0, 1}^(*k*−1)*×T*^ stack the column masks as rows. Define

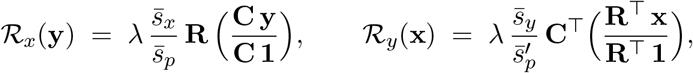

where the division inside each parenthesis is element–wise, **C 1** and **R**^⊤^**1** are the per–clump column and row sizes, *λ* is the repulsion strength, and 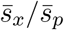 (resp.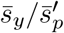) is a per-step rescaling that makes the mean magnitude of the penalty match the mean magnitude of the un-penalized score **xp** (resp. **yp**):

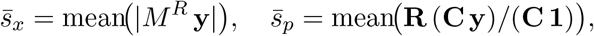

and analogously for the column side. Consider the row update. We first ask how many of the columns **y** in the current clump that each of the old clumps contains. This is what **C y** counts. Dividing it by the size *C***1** of the old clumps turns it into the fraction of the current clump’s columns that each old clump contains. We then transport this back to rows through *R*: a row is penalized in proportion to how strongly it participates in previous clumps whose columns overlap with the current clump candidate. In short, the penalty on row *i* is the sum over old clumps that contain row *i*, weighted by how much each of those old clumps’ columns overlaps with the current **y**. The column update is symmetric. The dynamic rescaling decouples the choice of *λ* from the absolute scale of *M* ^*R*^ and from how many clumps have already been found.

#### 4.2.3 Containment overlap and DBSCAN clustering of many clumps

To get a collection of *K* clumps for further analyses, we call the single-clump procedure *K* times in sequence, growing **R** and **C** by one column and one row, respectively, after each call. The first clump runs with ℛ_*x*_ = ℛ_*y*_ = 0 and every subsequent call sees a non-trivial repulsion. The resulting collection of *K* clumps will typically contain many overlapping clumps. This structure can be used to infer the geometry of the covariates we are trying to infer. This is done by clustering the containment overlap between the clump submatrices.

For clumps *k* and *ℓ*, the rectangular intersection **x**^(*k*)^ ∩ **x**^(*ℓ*)^ on rows and **y**^(*k*)^ ∩ **y**^(*ℓ*)^ on columns has size

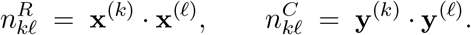

The product 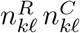 is the area of the rectangular intersection viewed as a submatrix of *M*. Normalize by the smaller of the two rectangle areas |**x**^(*k*)^| |**y**^(*ℓ*)^| and |**x**^(*ℓ*)^| |**y**^(*k*)^| to obtain a containment score,

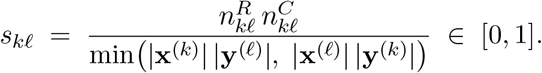

We have *s*_*kℓ*_ = 1 when one clump’s rectangle is fully contained in the other’s, and *s*_*kℓ*_ = 0 when they are disjoint on at least one side. Thus, it is not enough for clumps to overlap in only their selected rows or columns; they must overlap in both. The corresponding distance *d*_*kℓ*_ = 1 − *s*_*kℓ*_ is used as a precomputed metric for DBSCAN clustering with parameters *ε* and minPts. Noise points (DBSCAN label −1) are discarded. Note that the containment overlap distance is not a metric in the mathematical sense (the triangle inequality may fail). However, it fits our intuition well and we have found that it works well in practice, as DBSCAN does not assume a metric, only a symmetric pairwise distance.

#### 4.2.4 MDS embeddings of clustered clumps

For visualization, we have chosen to display the different clump clusters together (to preserve space), but we do have access to the cluster labels. There are a couple of different matrices we could apply dimensionality reduction on (here, MDS), to visualize the clump-structure. We could let the clumps (columns) of the indicator matrix **R** be points, or the clumps (rows) of the of the indicator matrix **C**, or the clumps (rows or columns) of the containment overlap matrix *s*_*kℓ*_. If the clumps indeed are local, the two indicator matrices should show the same qualitative geometry. The embedding of the containment overlaps is the strictest, because both the rows and columns need to be similar for two clumps to be close. This is why we chose to use this embedding in Figure 4, 6, and 11.

#### 4.2.5 Finding structure in the motor cortex

##### The data matrix

The data matrix 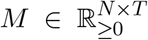 is made by binning the spikes into 10 ms bins, smoothing the neurons with a Gaussian kernel (*σ* = 100 ms), applying a 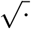 transform, and downsampling to every 5th bin.

##### Clumping and clustering

Like for the toy data, we make many clumps and cluster them with DBSCAN. The only difference is that, to make the clustering more robust, we expand each clump a bit around itself by running one fixed-point iteration with larger thresholds *t*_*r*_ and *t*_*c*_. For the motor cortex data, we used *t*_*r*_ = *t*_*c*_ = 0.25. Note that this results is substantially smaller clumps than if we had used these larger thresholds for the entire optimization. We also apply this expansion when visualizing the clump embeddings to make them look less scattered.

#### 4.2.6 Hyperparameters used in the paper

The clump figures in this paper use the following settings.

| Symbol | Meaning | Continuous<br>toy data | Discrete<br>toy data | Motor<br>cortex |
| --- | --- | --- | --- | --- |
| $t_r$ | row threshold (stds above mean) | 1.0 | 0.6 | 1.5 |
| $t_c$ | column threshold (stds above mean) | 1.0 | 0.6 | 1.5 |
| $K$ | number of clumps mined | 1000 | 1000 | 1000 |
| $\lambda$ | repulsion strength | 0.01 | 0.0 | 0.1 |
| $\tau_{\text{conv}}$ | cosine-distance convergence tolerance | 0.01 | 0.01 | 0.01 |
| $S_{\text{max}}$ | max iterations per clump (then restart) | 1000 | 1000 | 1000 |
| $\varepsilon$ | DBSCAN radius on $d = 1 - s$ | 0.4 | 0.2 | 0.5 |
| minPts | DBSCAN minimum cluster size | 10 | 25 | 15 |

## Appendix Related ideas

The quest to disentangle data into its underlying interpretable composites has long been ongoing. Many fields encounter problems that are similar to those we discuss here in the context of neural data analysis and many methods have been proposed to overcome those challenges. We hope that the principle of bidirectional locality introduced in this paper can provide a novel perspective on many of these works. Naturally, our discussion in this section will be far from complete. But we hope to give at least a rough sketch of the landscape.

- **Topological data analysis and Dowker duality**. Although we never make it explicit, this work is heavily inspired by a classical result from the mathematical field of topology under the name of *Dowker duality*^56^. This result establishes that the Dowker row complex *R*(*A*) and the Dowker column complex *C*(*A*) derived from a binary relation *A* are *always homotopy equivalent*. To be more precise, we should say that this work is inspired by a particularly simple proof strategy for Dowker duality^57^, where rows index a good open cover of the column complex. The proof is then reduced to a simple application of the *nerve lemma* to this open cover. The same strategy can be applied the other way around to a good open cover of the row complex indexed by the columns of *A*. This construction motivated us to study the intrinsic fields of rows and columns in the present paper. Our intuition is, that when the intrinsic fields are localized, a similar logic as in the proof of Dowker duality applies and the respective row and column spaces are dual. Crucially, and this is different from the topological Dowker duality, the locality condition on the intrinsic fields does not hold automatically. We saw that such failure can indeed coincide with the duality of row and column representations of the data breaking down. Our intuition is that in a bidirectionally local situation a metric form of Dowker duality holds and thus both the row and column representations hold the same self-coherent information. This intuition, as discussed here, is obviously informal and it is outside the theoretical scope and notation established here to provide more formal arguments. In^40^ we formally prove that bidirectionally locality of rows and columns of a data matrix *X* yields a bound on the interleaving between the Vietoris Rips filtrations of *X* and *X*^*T*^. This can be interpreted as a metric approximation to Dowker duality. We are currently working on more mathematical expositions and results around this.
- **Biclustering**. A clump, as we define it in this paper, is a *bicluster* : a dense submatrix defined by a jointly selected set of rows and columns. The idea of mining dense submatrices goes back to Good^26^, where he also uses the word clump that we adopted here. The fixed-point mining iterations we use are almost verbatim the signature algorithm of^41^, part of a large biclustering literature in gene-expression analysis^46;47;58^. We put the method into a new context and argue in (Section 4.2.1) that under additive mixed selectivity a dense submatrix is, with overwhelming probability, local on a single covariate, so clumps become point-like proxies that tile the latent; we then cluster and embed the clumps to recover covariate geometry, a step the expression-data literature does not take. As an important technical contribution we add a *repulsion term* to the algorithm. We found this to be crucial for covering a continuous covariate with clumps mined through the iterative signature algorithm, which without the term always converged to a few non-overlapping regions on the space.
- **Formal concept analysis**. Formal concept analysis (FCA) studies a binary relation between a set of objects and a set of attributes — a *formal context* — through its *formal concepts*^59;60^. A formal concept is a *maximal all-ones submatrix* of the relation. This is the binary special case of our clumps: a clump is the density-based, noise-tolerant relaxation of a formal concept, defined on the real-valued matrix *M* rather than a binary relation and requiring the sub-block to be dense rather than strictly all-ones. Where FCA enumerates the full concept lattice exactly, we instead collect a diverse bank of approximate concepts and encourage coverage of the underlying covariate through the repulsion term. We experimented a lot with different ways of binarizing a neural activity matrix and subsampling (approximate) maximal bicliques of the resulting row-column bipartite graph. These experiments had some success but ultimately fell victim to the combinatorial explosion of possible bicliques in dense regions.
- **Community detection**. Finding clusters or communities in graphs is known as community detection^49^. In our case, the graph could be an adjacency matrix between neurons or timebins, or even a bipartite graph between neurons are timebins. Communities are typically defined as being assortative, with a connection within a community being more likely than a connection between communities. However, different notions exist^61^, leading to a wide variety of methods, such as^62^. When (assortative) communities are allowed to overlap, the problem starts looking like separating covariates in the discrete situation discussed here, as every node might be “tuned” to multiple “covariates”. One method developed to find overlapping communities^50^ is based on another idea closely related to our work, nonnegative matrix factorization, which we discuss next.
- **Nonnegative matrix factorization**. In Nonnegative Matrix Factorization (NMF) the data matrix is typically factorized into two matrices, *M* ≈ *WH*^23;63^. The factors learned by NMF are often described as parts-based and interpretable, due to the non-negativity constraint prohibiting complex cancellations of positive and negative values. This was one of the first methods we tried for separating covariates in mixed selectivity data. Our intuition is that NMF would find structure that’s shared by the row- and column-space of a matrix. One loose justification for this is that neurons are recovered by adding together columns of *W* (neurons “parts”) according to rows in *H* and time bins are recovered by adding together rows of *H* (timebin “parts”) according to columns of *W*. Other reasons why NMF seemed promising was its close relation to spectral clustering and biclustering of a bipartite graph^64;65^ and the suggestions that it could recover a low-dimensional manifold embedded in high-dimensional space^66^. While (a variation of) NMF worked well in certain simulated settings, it was very sensitive to the sampling density of the latent covariates. This is because its only objective is to reconstruct the data. If and when it recovered the latent covariates, it was merely a by-product of this objective. This motivated us to develop methods more explicitly optimizing for the bidirectional locality we actually wanted. We also want to note that while coherent projections and clumps don’t optimize for matrix reconstruction, it’s not difficult get reconstruction estimates from them. For coherent projections, we can regress the neurons on the modes of the projected matrix *E*. For clumps, we can fit a standard NMF where the stacked clumps constrain and mask *W* and *H*.
- **Dimensionality reduction and disentanglement**. Dimensionality reduction and disentanglement have the shared goal of finding a compressed description of the data. Disentanglement has the additional objective of decomposing the data into separate sources of variation. Sometimes, when the data contains several “pure” populations (the second example in Section 2.2), dimensionality reduction can seem to do this by making a disjoint union. However, this is rarely the case. More often, data is mixed in both rows and columns, which unsupervised dimensionality reduction methods such as PCA, ICA, Isomap, and UMAP^28;52;67;68^ reduce along a single direction and therefore cannot disentangle. A second group of methods supplies structure in the missing direction from outside the matrix: dPCA and nonlinear ICA^21;22;69;70^ condition on labels or an auxiliary variable, without which the nonlinear problem is ill-defined^71^. Tensor methods, in contrast, are unsupervised and use every direction at once, but they require a neuron × time × trial tensor. TCA^72^ fits components of tensor rank one, an outer product of three vectors that fix a single neuron-, trial-, and time-pattern. sliceTCA^73^ relaxes this to slice rank one, an outer product of one vector and one unconstrained matrix, so that each component is constrained along a single axis and free across the other two. These constraints don’t enforce locality, so the components are not local on any covariate and cannot serve as points from which to recover its geometry. We instead present bidirectional locality as a signature of a disentangled representation, and show how it identifies such representations, without trials or labels.
- **Abstract representations**. One line of work argue brains and machines uses the same representations of variables that are used in different contexts^9;19;74^. These variables are thus represented in an abstract (or disentangled) format, because it allows for generalization or abstraction over tasks. Abstract representations can be identified by decoding the information/variable with a decoder trained on one task, during another task. Notably, this holds despite neurons exhibiting mixed selectivity to the relevant task variables^74^. We already hinted towards this work in the introduction. Here, we make the connection more explicit. First, abstract representations are typically low-dimensional, like those of the covariates we have studied here. Second, the argument for abstract representations is similar to the argument for recovering covariates. That is, instead of representing every experience or task as its own high-dimensional thing (in the product space), it’s more compact and efficient to decompose it into multiple low-dimensional variables. Third, the prevalence of abstract representations strongly supports our suspicion that neural recordings might entangle different covariates, highlighting the need for methods that can separate them.
- **Field-like coding in machine learning**. When studying how concepts are represented in artificial neural networks, such as large language models, the concepts have often been equated with linear directions. This is known as the linear representation hypothesis^53;54^. However, features can also employ the field-like and manifold-tiling coding we see in neuroscience^75;76^. One recent advance is the use of such field-like coding to interpret large language^5^ and vision^77^ models. While this work often relies on sparsity to generate the field-like coding, we, in contrast, optimize^40^ and look for it explicitly. Finally, we believe that linear and field-like coding are two qualitatively different strategies and that their relative strengths and weaknesses is a promising future direction.
- **Disentanglement in machine learning**. Unsupervised disentanglement has been pursued with variational and adversarial objectives^25;78^ and formalized definitionally^79^. A key negative result is that disentanglement is impossible without inductive biases or weak supervision^80^; subsequent work supplies such biases through temporal sparsity or subspace structure^81;82^. Our inductive bias is bidirectional locality together with nonnegativity, and our “generative factors” are the covariates an animal experiences. We contribute both a diagnostic (does a representation carry the signature?) and explicit constructions, rather than a single generative model.

## Acknowledgments

The work was supported by a grant from the Research Council of Norway (iMOD, NFR grant 325114); a Centre of Excellence grant (Centre for Algorithms in the Cortex, grant 332640) from the Research Council of Norway; and the Department of Mathematics at NTNU.

## References

[1] H Sebastian Seung and Daniel D Lee. The manifold ways of perception. science, 290(5500): 2268–2269, 2000.

[2] Yoshua Bengio, Aaron Courville, and Pascal Vincent. Representation learning: A review and new perspectives. IEEE Transactions on Pattern Analysis and Machine Intelligence, 35(8): 1798–1828, 2013. doi: 10.1109/TPAMI.2013.50.

[3] Charles Fefferman, Sanjoy Mitter, and Hariharan Narayanan. Testing the manifold hypothesis. Journal of the American Mathematical Society, 29(4):983–1049, 2016. doi: 10.1090/jams/852.

[4] Juan A. Gallego, Matthew G. Perich, Lee E. Miller, and Sara A. Solla. Neural manifolds for the control of movement. Neuron, 94(5):978–984, 2017. doi: 10.1016/j.neuron.2017.05.025.

[5] Usha Bhalla, Thomas Fel, Can Rager, Sheridan Feucht, Tal Haklay, Daniel Wurgaft, Siddharth Boppana, Matthew Kowal, Vasudev Shyam, Jack Merullo, et al. Do sparse autoencoders capture concept manifolds? arXiv preprint arXiv:2604.28119, 2026.

[6] SueYeon Chung and L. F. Abbott. Neural population geometry: An approach for understanding biological and artificial neural networks. Current Opinion in Neurobiology, 70:137–144, 2021. doi: 10.1016/j.conb.2021.10.010.

[7] Irina Higgins, L. Chang, Victoria Langston, Demis Hassabis, Christopher Summerfield, Doris Tsao, and Matthew Botvinick. Unsupervised deep learning identifies semantic disentanglement in single inferotemporal face patch neurons. Nature communications, 12(1):6456, 2021.

[8] W. Jeffrey Johnston and Stefano Fusi. Abstract representations emerge naturally in neural networks trained to perform multiple tasks. Nature Communications, 14(1):1040, 2023.

[9] James C. R. Whittington, Will Dorrell, Surya Ganguli, and Timothy E. J. Behrens. Disentanglement with biological constraints: A theory of functional cell types. In International Conference on Learning Representations (ICLR), 2023. arXiv:2210.01768.

[10] Adam Shai, Loren Amdahl-Culleton, Casper L Christensen, Henry R Bigelow, Fernando E Rosas, Alexander B Boyd, Eric A Alt, Kyle J Ray, and Paul M Riechers. Transformers learn factored representations. arXiv preprint arXiv:2602.02385, 2026.

[11] Mattia Rigotti, Omri Barak, Melissa R. Warden, Xiao-Jing Wang, Nathaniel D. Daw, Earl K. Miller, and Stefano Fusi. The importance of mixed selectivity in complex cognitive tasks. Nature, 497(7451):585–590, 2013.

[12] Francesca Sargolini, Marianne Fyhn, Torkel Hafting, Bruce L McNaughton, Menno P Witter, May-Britt Moser, and Edvard I Moser. Conjunctive representation of position, direction, and velocity in entorhinal cortex. Science, 312(5774):758–762, 2006.

[13] Mehrdad Kashefi, Jonathan A Micheals, Rhonda Kersten, Jonathan C Lau, Jörn Diedrichsen, and J Andrew Pruszynski. Compositional neural dynamics during reaching. bioRxiv, pages 2025–09, 2025.

[14] Debora Ledergerber, Claudia Battistin, Jan Sigurd Blackstad, Richard J Gardner, Menno P Witter, May-Britt Moser, Yasser Roudi, and Edvard I Moser. Task-dependent mixed selectivity in the subiculum. Cell reports, 35(8), 2021.

[15] Shinichiro Kira, Houman Safaai, Ari S Morcos, Stefano Panzeri, and Christopher D Harvey. A distributed and efficient population code of mixed selectivity neurons for flexible navigation decisions. Nature communications, 14(1):2121, 2023.

[16] Kay M Tye, Earl K Miller, Felix H Taschbach, Marcus K Benna, Mattia Rigotti, and Stefano Fusi. Mixed selectivity: Cellular computations for complexity. Neuron, 112(14):2289–2303, 2024.

[17] Matthew T Kaufman, Marcus K Benna, Mattia Rigotti, Fabio Stefanini, Stefano Fusi, and Anne K Churchland. The implications of categorical and category-free mixed selectivity on representational geometries. Current opinion in neurobiology, 77:102644, 2022.

[18] Stefano Fusi, Earl K. Miller, and Mattia Rigotti. Why neurons mix: high dimensionality for higher cognition. Current Opinion in Neurobiology, 37:66–74, 2016.

[19] W. Jeffrey Johnston, Stephanie E. Palmer, and David J. Freedman. Nonlinear mixed selectivity supports reliable neural computation. PLOS Computational Biology, 16(2):e1007544, 2020.

[20] Corey J Maley. Analog and digital, continuous and discrete. Philosophical Studies, 155(1): 117–131, 2011.

[21] Dmitry Kobak, Wieland Brendel, Christos Constantinidis, Claudia E. Feierstein, Adam Kepecs, Zachary F. Mainen, Ranulfo Romo, Xue-Lian Qi, Naoshige Uchida, and Christian K. Machens. Demixed principal component analysis of neural population data. eLife, 5:e10989, 2016.

[22] Steffen Schneider, Jin Hwa Lee, and Mackenzie Weygandt Mathis. Learnable latent embeddings for joint behavioural and neural analysis. Nature, 617(7960):360–368, 2023.

[23] Te-Won Lee. Independent component analysis. In Independent component analysis: Theory and applications, pages 27–66. Springer, 1998.

[24] Emily L Mackevicius, Andrew H Bahle, Alex H Williams, Shijie Gu, Natalia I Denisenko, Mark S Goldman, and Michale S Fee. Unsupervised discovery of temporal sequences in high-dimensional datasets, with applications to neuroscience. Elife, 8:e38471, 2019.

[25] Irina Higgins, Loic Matthey, Arka Pal, Christopher Burgess, Xavier Glorot, Matthew Botvinick, Shakir Mohamed, and Alexander Lerchner. beta-VAE: Learning basic visual concepts with a constrained variational framework. In International Conference on Learning Representations (ICLR), 2017.

[26] I. J. Good. The Estimation of Probabilities: An Essay on Modern Bayesian Methods. Research Monograph No. 30. MIT Press, Cambridge, MA, 1965. Discusses “clumps” as dense submatrices.

[27] Apostolos P. Georgopoulos, Andrew B. Schwartz, and Ronald E. Kettner. Neuronal population coding of movement direction. Science, 233(4771):1416–1419, 1986.

[28] John P. Cunningham and Byron M. Yu. Dimensionality reduction for large-scale neural recordings. Nature Neuroscience, 17(11):1500–1509, 2014.

[29] Saurabh Vyas, Matthew D. Golub, David Sussillo, and Krishna V. Shenoy. Computation through neural population dynamics. Annual Review of Neuroscience, 43:249–275, 2020.

[30] Carina Curto and Vladimir Itskov. Cell groups reveal structure of stimulus space. PLoS Computational Biology, 4(10):e1000205, 2008.

[31] Chad Giusti, Eva Pastalkova, Carina Curto, and Vladimir Itskov. Clique topology reveals intrinsic geometric structure in neural correlations. Proceedings of the National Academy of Sciences, 112(44):13455–13460, 2015.

[32] Rishidev Chaudhuri, Berk Gerçek, Biraj Pandey, Adrien Peyrache, and Ila Fiete. The intrinsic attractor manifold and population dynamics of a canonical cognitive circuit across waking and sleep. Nature Neuroscience, 22(9):1512–1520, 2019.

[33] Nikolaus Kriegeskorte, Marieke Mur, and Peter A. Bandettini. Representational similarity analysis – connecting the branches of systems neuroscience. Frontiers in Systems Neuroscience, 2:4, 2008.

[34] Ann E. Sizemore, Jennifer E. Phillips-Cremins, Robert Ghrist, and Danielle S. Bassett. The importance of the whole: topological data analysis for the network neuroscientist. Network Neuroscience, 3(3):656–673, 2019.

[35] Arianna Di Bernardo, Adrian Valente, Francesca Mastrogiuseppe, and Srdjan Ostojic. Shaping manifolds in equivariant recurrent neural networks. arXiv preprint arXiv:2511.04802, 2025.

[36] Matthew T Kaufman, Mark M Churchland, Stephen I Ryu, and Krishna V Shenoy. Cortical activity in the null space: permitting preparation without movement. Nature neuroscience, 17 (3):440–448, 2014.

[37] Matthew R Ginther, Devin F Walsh, and Seth J Ramus. Hippocampal neurons encode different episodes in an overlapping sequence of odors task. Journal of Neuroscience, 31(7):2706–2711, 2011.

[38] Richard J. Gardner, Erik Hermansen, Marius Pachitariu, Yoram Burak, Nils A. Baas, Benjamin A. Dunn, May-Britt Moser, and Edvard I. Moser. Toroidal topology of population activity in grid cells. Nature, 602(7895):123–128, 2022.

[39] Erik Hermansen, David A Klindt, and Benjamin A Dunn. Uncovering 2-d toroidal representations in grid cell ensemble activity during 1-d behavior. Nature Communications, 15(1):5429, 2024.

[40] Sigurd Gaukstad, Melvin Vaupel, Valdemar Kargård Olsen, Erik Hermansen, and Benjamin Adric Dunn. Learning coherent representations: A topological approach to interpretability. In Forty-third International Conference on Machine Learning, 2026. URL https://openreview.net/forum?id=VnlpOuA3Bs.

[41] Sven Bergmann, Jan Ihmels, and Naama Barkai. Iterative signature algorithm for the analysis of large-scale gene expression data. Physical review E, 67(3):031902, 2003.

[42] Hanne Stensola, Tor Stensola, Trygve Solstad, Kristian Frøland, May-Britt Moser, and Edvard I. Moser. The entorhinal grid map is discretized. Nature, 492(7427):72–78, 2012.

[43] Bartul Mimica, Tuce Tombaz, Claudia Battistin, Jingyi Guo Fuglstad, Benjamin A. Dunn, and Jonathan R. Whitlock. Behavioral decomposition reveals rich encoding structure employed across neocortex in rats. Nature Communications, 14(1):3947, 2023.

[44] David M Blei, Andrew Y Ng, and Michael I Jordan. Latent dirichlet allocation. Journal of machine Learning research, 3(Jan):993–1022, 2003.

[45] Thomas Hofmann. Probabilistic latent semantic analysis. arXiv preprint arXiv:1301.6705, 2013.

[46] Sara C. Madeira and Arlindo L. Oliveira. Biclustering algorithms for biological data analysis: a survey. IEEE/ACM Transactions on Computational Biology and Bioinformatics, 1(1):24–45, 2004.

[47] Yizong Cheng and George M. Church. Biclustering of expression data. In Proceedings of the International Conference on Intelligent Systems for Molecular Biology (ISMB), volume 8, pages 93–103, 2000.

[48] Dhruva Karkada, James Simon, Yasaman Bahri, and Michael Deweese. Closed-form training dynamics reveal learned features and linear structure in word2vec-like models. Advances in Neural Information Processing Systems, 38:3967–4000, 2026.

[49] Santo Fortunato and Darko Hric. Community detection in networks: A user guide. Physics reports, 659:1–44, 2016.

[50] Jaewon Yang and Jure Leskovec. Overlapping community detection at scale: a nonnegative matrix factorization approach. In Proceedings of the sixth ACM international conference on Web search and data mining, pages 587–596, 2013.

[51] Daniel D. Lee and H. Sebastian Seung. Learning the parts of objects by non-negative matrix factorization. Nature, 401(6755):788–791, 1999.

[52] Aapo Hyvärinen and Erkki Oja. Independent component analysis: algorithms and applications. Neural Networks, 13(4–5):411–430, 2000.

[53] Kiho Park, Yo Joong Choe, and Victor Veitch. The linear representation hypothesis and the geometry of large language models. arXiv preprint arXiv:2311.03658, 2023.

[54] Nelson Elhage, Tristan Hume, Catherine Olsson, Nicholas Schiefer, Tom Henighan, Shauna Kravec, Zac Hatfield-Dodds, Robert Lasenby, Dawn Drain, Carol Chen, et al. Toy models of superposition. arXiv preprint arXiv:2209.10652, 2022.

[55] Sigurd Gaukstad. Topological Approaches for Discovering and Learning Neural Representations. PhD thesis, Norges teknisk-naturvitenskapelige universitet, 2026. URL https://api.nva.unit.no/publication/019e88236434-543789d6-940c-4428-9115-99b35940e650. Nva type: DegreePhd.

[56] C. H. Dowker. Homology groups of relations. Annals of Mathematics, 56(1):84–95, 1952.

[57] Anders Björner. Topological methods. Handbook of combinatorics, 2:1819–1872, 1995.

[58] Victor A Padilha and Ricardo JGB Campello. A systematic comparative evaluation of biclustering techniques. BMC bioinformatics, 18(1):55, 2017.

[59] Rudolf Wille. Restructuring lattice theory: an approach based on hierarchies of concepts. In Ivan Rival, editor, Ordered Sets, volume 83 of NATO Advanced Study Institutes Series, pages 445–470. Reidel, Dordrecht, 1982.

[60] Bernhard Ganter and Rudolf Wille. Formal Concept Analysis: Mathematical Foundations. Springer, Berlin, Heidelberg, 1999.

[61] Michael T Schaub, Jean-Charles Delvenne, Martin Rosvall, and Renaud Lambiotte. The many facets of community detection in complex networks. Applied network science, 2(1):4, 2017.

[62] Tiago P Peixoto. Bayesian stochastic blockmodeling. Advances in network clustering and blockmodeling, pages 289–332, 2019.

[63] Patrik O Hoyer. Non-negative matrix factorization with sparseness constraints. Journal of machine learning research, 5(Nov):1457–1469, 2004.

[64] Chris Ding, Xiaofeng He, and Horst D Simon. On the equivalence of nonnegative matrix factorization and spectral clustering. In Proceedings of the 2005 SIAM international conference on data mining, pages 606–610. SIAM, 2005.

[65] Chris Ding, Tao Li, Wei Peng, and Haesun Park. Orthogonal nonnegative matrix t-factorizations for clustering. In Proceedings of the 12th ACM SIGKDD international conference on Knowledge discovery and data mining, pages 126–135, 2006.

[66] Deng Cai, Xiaofei He, Jiawei Han, and Thomas S Huang. Graph regularized nonnegative matrix factorization for data representation. IEEE transactions on pattern analysis and machine intelligence, 33(8):1548–1560, 2010.

[67] Joshua B Tenenbaum, Vin de Silva, and John C Langford. A global geometric framework for nonlinear dimensionality reduction. science, 290(5500):2319–2323, 2000.

[68] Leland McInnes, John Healy, and James Melville. Umap: Uniform manifold approximation and projection for dimension reduction. arXiv preprint arXiv:1802.03426, 2018.

[69] Aapo Hyvärinen, Hiroaki Sasaki, and Richard E. Turner. Nonlinear ICA using auxiliary variables and generalized contrastive learning. In Proceedings of the 22nd International Conference on Artificial Intelligence and Statistics (AISTATS), pages 859–868, 2019.

[70] Ding Zhou and Xue-Xin Wei. Learning identifiable and interpretable latent models of high-dimensional neural activity using pi-vae. Advances in neural information processing systems, 33:7234–7247, 2020.

[71] Aapo Hyvärinen and Petteri Pajunen. Nonlinear independent component analysis: Existence and uniqueness results. Neural Networks, 12(3):429–439, 1999.

[72] Alex H Williams, Tony Hyun Kim, Forea Wang, Saurabh Vyas, Stephen I Ryu, Krishna V Shenoy, Mark Schnitzer, Tamara G Kolda, and Surya Ganguli. Unsupervised discovery of demixed, low-dimensional neural dynamics across multiple timescales through tensor component analysis. Neuron, 98(6):1099–1115, 2018.

[73] Arthur Pellegrino, Heike Stein, and N Alex Cayco-Gajic. Dimensionality reduction beyond neural subspaces with slice tensor component analysis. Nature Neuroscience, 27(6):1199–1210, 2024.

[74] Silvia Bernardi, Marcus K. Benna, Mattia Rigotti, Jérôme Munuera, Stefano Fusi, and C. Daniel Salzman. The geometry of abstraction in the hippocampus and prefrontal cortex. Cell, 183(4):954–967, 2020.

[75] Bruno A Olshausen and David J Field. Emergence of simple-cell receptive field properties by learning a sparse code for natural images. Nature, 381(6583):607–609, 1996.

[76] Anirvan M. Sengupta, Mariano Tepper, Cengiz Pehlevan, Alexander Genkin, and Dmitri B. Chklovskii. Manifold-tiling localized receptive fields are optimal in similarity-preserving neural networks. In Advances in Neural Information Processing Systems (NeurIPS), pages 7080–7090, 2018.

[77] Thomas Fel, Matthew Kowal, Mozes Jacobs, Dron Hazra, Usha Bhalla, Lee Sharkey, Lucius Bushnaq, Satchel Grant, Tal Haklay, Thomas Icard, et al. Structuring sparsity: Block-sparse featurizers capture visual concept manifolds. arXiv preprint arXiv:2606.25234, 2026.

[78] Xi Chen, Yan Duan, Rein Houthooft, John Schulman, Ilya Sutskever, and Pieter Abbeel. InfoGAN: Interpretable representation learning by information maximizing generative adversarial nets. In Advances in Neural Information Processing Systems (NeurIPS), pages 2172–2180, 2016.

[79] Irina Higgins, David Amos, David Pfau, Sebastien Racaniere, Loic Matthey, Danilo Rezende, and Alexander Lerchner. Towards a definition of disentangled representations. arXiv preprint arXiv:1812.02230, 2018.

[80] Francesco Locatello, Stefan Bauer, Mario Lucic, Gunnar Rätsch, Sylvain Gelly, Bernhard Schölkopf, and Olivier Bachem. Challenging common assumptions in the unsupervised learning of disentangled representations. In Proceedings of the 36th International Conference on Machine Learning (ICML), pages 4114–4124, 2019.

[81] David A. Klindt, Lukas Schott, Yash Sharma, Ivan Ustyuzhaninov, Wieland Brendel, Matthias Bethge, and Dylan Paiton. Towards nonlinear disentanglement in natural data with temporal sparse coding. In International Conference on Learning Representations (ICLR), 2021. arXiv:2007.10930.

[82] David Pfau, Irina Higgins, Aleksandar Botev, and Sébastien Racanière. Disentangling by subspace diffusion. In Advances in Neural Information Processing Systems (NeurIPS), pages 17403–17415, 2020.

